# Kappa but not delta or mu opioid receptors form homodimers at low membrane densities

**DOI:** 10.1101/2020.03.22.002394

**Authors:** Kristina Cechova, Chenyang Lan, Matus Macik, Nicolas P. F. Barthes, Manfred Jung, Maximilian H. Ulbrich

**Affiliations:** Department of Biomathematics, Institute of Physiology, Czech Academy of Sciences; Department of Biochemistry, Faculty of Science, Charles University; Faculty of Biology, University of Freiburg; Independet Scholar; Institute of Pharmaceutical Sciences, University of Freiburg; CIBSS – Centre for Integrative Biological Signalling Studies, University of Freiburg; Institute of Internal Medicine IV, Medical Center of the University of Freiburg; BIOSS Centre for Biological Signalling Studies, University of Freiburg

## Abstract

Opioid receptors (ORs) have been observed as homo- and heterodimers, but it is unclear if the dimers are stable under physiological conditions, and whether monomers or dimers comprise the predominant fraction in a cell. Here we use three live-cell imaging approaches to assess dimerization of ORs at different expression levels. At high membrane densities, a split GFP assay reveals that κOR dimerizes, while μOR and δOR stay monomeric. In contrast, single-molecule imaging showed no κOR dimers at low receptor densities. To reconcile our seemingly contradictory results, we used a high-density single-molecule assay to assess membrane protein interactions at densities up to 100x higher than conventional single-molecule imaging. We observe that κOR is monomeric at low densities and forms dimers at densities that are considered physiological. In contrast, μOR and δOR stay monomeric even at the highest densities covered by our approach. The observation of long-lasting κOR dimers but not higher-order aggregates suggests that ORs dimerize through a single, specific interface.

## Introduction

ORs are G protein-coupled receptors (GPCRs) from class A with three genes coding for the δOR, κOR, and μOR. The μOR is the most prominent clinical target for pain medication. However, side effects of opiates like respiratory depression and the potential for opioid addiction have increased the efforts to develop drugs with novel pharmacological profiles. Also, it has become clear in recent years that differential control of downstream mechanisms (G protein activation vs. β- arrestin recruitment) or restriction of drugs to the peripheral nervous system allows to further reduce unwanted side effects. Therefore, improving our understanding of OR activation and signaling will support the development of novel treatments.

Based on functional characterization of ORs, more subtypes than δ, κ, and μ were proposed, which can be explained by the existence of splicing variants, posttranslational modifications and/or direct interactions between receptors. Dimerization of ORs has been covered in multiple studies, yet the conclusions were contradictory, as for many other GPCRs, likely due to the use of differing methodological approaches and experimental conditions [1]. For all three ORs, dimerization has been observed [2–6]. So far, the major techniques to assess dimerization of ORs were co-immunoprecipitation followed by Western blotting, and bioluminescence resonance energy transfer (BRET). Both are bulk techniques, where the signal is obtained from a large population of cells. Not only is the major part of the signal caused by a small, highly expressing fraction of the cells, but in addition, these few cells express the receptors at the highest density. Therefore, the signal mainly reflects the receptor’s behavior at a membrane density that is far above the physiological range, and cannot capture the state of the receptors at low membrane densities, as they prevail in many cells *in vivo*.

To assess specific interactions of membrane proteins that are likely to exist *in vivo* as well, single-molecule imaging in living cells is the perfect tool. At densities around 0.1-1 / µm^2^, single fluorescently labeled molecules can be tracked, and dimerization or higher-order cluster formation can be observed as co-localization of differentially labeled receptors or can be deduced from the analysis of intensity histograms from many individual receptor complexes [7, 8]. For several class A GPCRs, single-molecule imaging suggested the existence of dimers [7, 9, 10]. μOR seems to be monomeric when unliganded, but switches to a dimeric state when the ligand DAMGO is bound, but interestingly stays monomeric upon binding of morphine [11, 12]. κOR has been observed to be primarily monomeric [13]. However, for these studies, experiments were performed at low densities < 1 / µm^2^. A recent study using single-molecule FRET also observed unliganded μOR to be monomeric at a density of up to 4 / µm^2^, and also at densities around 100 / µm^2^ using pulsed-interleaved excitation fluorescence cross-correlation spectroscopy (PIE-FCCS) [14]. No study so far investigated dimerization of the δOR at low densities. The observation of OR monomers at low densities and dimers in bulk experiments suggest that the dimerization depends on the receptor density in the membrane.

In this work, we set out to investigate OR dimerization at densities that were not covered previously, with the goal to determine the density at which a switch from monomers to dimers happens. We therefore quantitatively measured OR dimerization using three microscopy approaches that work at different densities. Our starting point was a split GFP fluorescence complementation assay in cells with expression levels in the range of 10–100 / µm^2^. This assay gave us a first indication that the κOR has a tendency to dimerize, while the δOR and μOR resembled the monomeric control at these densities, which was the PDGFRα transmembrane domain (PDGFRTM) [15]. However, with conventional dual-color single-molecule imaging at densities below 5 / µm^2^, we did not observe a significantly higher dimerization for κOR than for the other ORs. Therefore, we turned to a recently developed technique called PhotoGate that allows tracking of single molecules in an originally crowded environment by controlling the density of fluorescent molecules in a region of interest [16]. We quantified the monomer/dimer ratio at densities of up to 150 / µm^2^, a level that is thought to be in the physiological range for many GPCRs. δOR and μOR remained predominantly monomeric at all densities tested, whereas κOR formed dimers at higher levels, with a dissociation constant of *k*_*d*_ = 32 ± 15 / µm^2^.

## Results

### Split GFP complementation suggests a higher tendency of κOR to dimerize

To assess dimerization of ORs, we used fluorescence complementation of a split GFP. Here, two fragments of GFP, which on their own are not fluorescent, are fused to the target proteins. Upon encounter, the two fragments covalently bind to each other, which allows the GFP chromophore to form and fluorescence to recover [17]. In our case, we would co-transfect an OR fused to the first fragment with the same OR fused to the second fragment; in case of dimerization of the OR, fluorescence of the reconstituted GFP should appear.

One known disadvantage of the split GFP assay is that also non-interacting target proteins can lead to a certain degree of GFP complementation due to spontaneous encounters of the two fragments. Another problem is that fluorescence recovery is not proportional to the degree of dimer formation because once the two parts are connected, they cannot separate again. Both effects impede an accurate quantification of the dimerization process, and therefore the GFP complementation assay only allows a qualitative assessment of dimerization. Nevertheless, due to its simplicity, this assay has frequently been used [18].

Out of several different previously used split positions, we chose the one between the 10th and 11th beta strand of the GFP barrel, yielding one fragment containing the first ten strands (GFP10), and one fragment containing the last strand (GFP11) [19]. To determine the fraction of split GFP tags that recover fluorescence upon binding, we need to know the densities of the fragments and the density of recovered GFP. To this end, the two constructs carrying GFP10 and GFP11 were additionally tagged with mKate, which emits in the orange range, and the SNAP-tag, which we labeled with a far-red substrate (Fig. 1A). By comparing the intensities emitted from highly expressing cells with intensities of single molecules of GFP, mKate and the far-red labeled SNAP-tag, we calculated the absolute densities of these species in the membrane.

**Figure 1:**
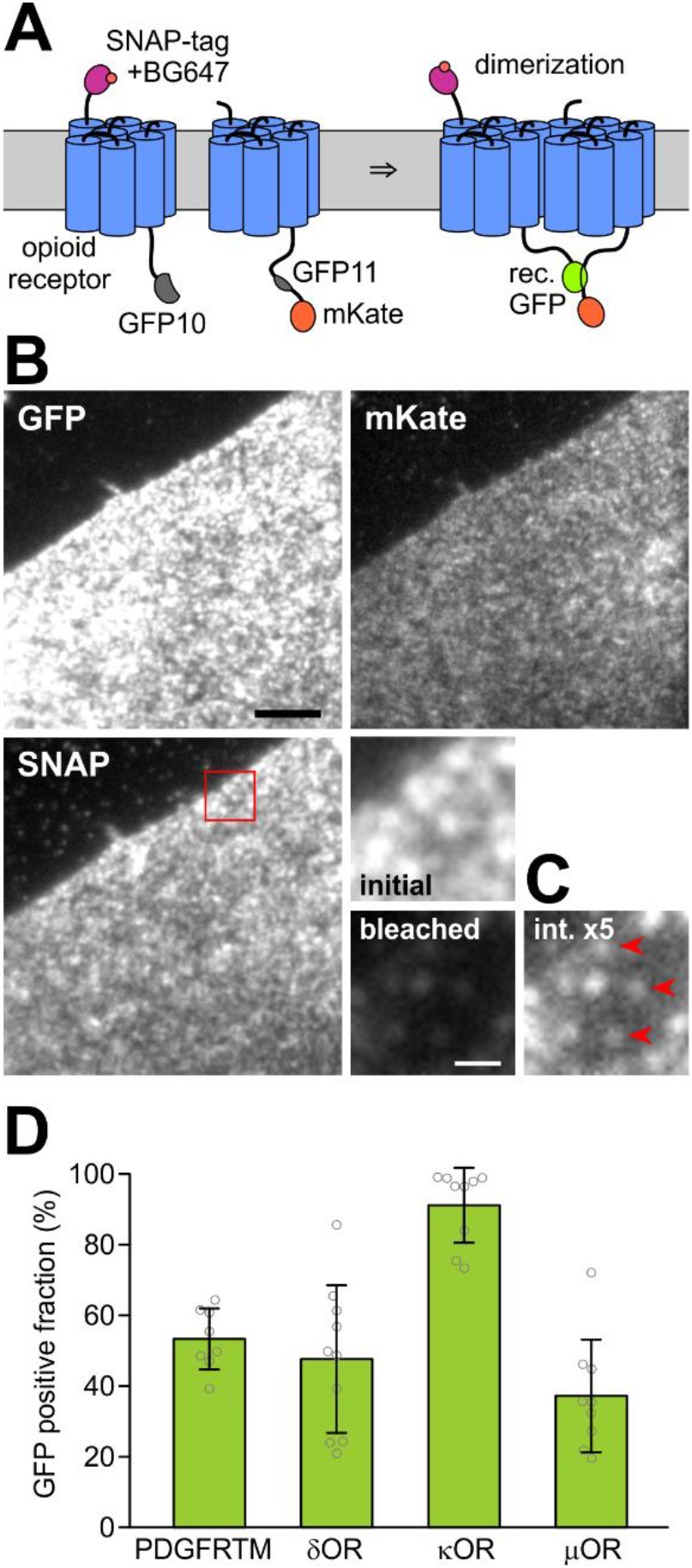
Split GFP assay for opioid receptor dimerization. **(A)** To determine the densities of the two differentially labeled OR subunits, one is labeled with a SNAP-tag and its far-red fluorescent substrate BG647, and the other with mKate which emits in the orange/red range. They are also labeled with two GFP fragments. Dimerization is detected by fluorescence recovery through complementation of the split GFP. **(B)** Images of a CHO cell expressing SNAP647-κOR-GFP10 and κOR-GFP11-mKate in the three channels. The red square marks the magnified area. Scale bar 5 µm. **(C)** Magnified area: Molecular densities were determined by measuring the initial fluorescence, bleaching the cell and then measuring the intensities of remaining single molecules. For the last image, the brightness is increased by a factor of 5. Red arrowheads mark single molecules. Scale bar 1 µm. **(D)** The GFP positive fraction for the ORs and the monomeric control PDGFRTM was calculated from apparent densities of SNAP647, mKate, and reconstituted GFP.

We generated pairs of δOR, κOR, or μOR, or the PDGFRTM as a negative control, fused with an N-terminal SNAP-tag and C-terminal GFP10, or C-terminal GFP11 and mKate, respectively, resulting in SNAP-X-GFP10 and X-GFP11-mKate, where X was δOR, κOR, μOR, or PDGFRTM (Fig. 1A). CHO cells expressing a matching pair were labeled with the far-red fluorescent dye Alexa Fluor 647 conjugated to benzylguanine (BG647), and imaged with 488 nm, 561 nm, and 637 nm excitation in Total Internal Reflection (TIR) configuration (Fig. 1B). From highly expressing cells (densities 7-90 / µm^2^, average 20-26 / µm^2^) with a similar density of mKate and BG647-labeled SNAP-tag (SNAP647), we recorded a snapshot for each wavelength range to determine the fluorescence intensity per area. For each of the three channels, single-molecule intensities were measured from nearly photobleached or low-expressing cells where individual, diffusing molecules were visible that photobleached in a single step (n ≥ 10 for each channel, Fig. 1C and Suppl. Fig. 1).

The molecule densities of SNAP647, mKate and (reconstituted) GFP tags were calculated from the ratios of intensities per area in the highly expressing cells to the intensities of single molecules from the low-density cells. The GFP-positive fraction was calculated as the ratio of GFP density to the smaller value of mKate and SNAP647 densities (for the rationale behind this approach, see Suppl. Note 1). For PDGFRTM, the GFP-positive fraction was 53 ± 9% (s.d., n = 8), and the smaller one of the mKate and SNAP647 surface density 23 ± 18 / µm^2^, for δOR 48 ± 21% (26 ± 23 / µm^2^, n = 10), for κOR 91 ± 11% (26 ± 22 / µm^2^, n = 9), and for the μOR 37 ± 16% (20 ± 14 / µm^2^, n = 9) (Fig. 1D).

The PDGFRTM, which served as a negative control since it is thought to be monomeric, also had a sizeable GFP-positive fraction due to reasons discussed above. The significantly larger GFP-positive fraction (p < 0.0007) for the κOR than for the negative control PDGFRTM, δOR, and μOR suggests that the κOR has a higher tendency to dimerize than the other two ORs. The GFP-positive fraction for δOR and μOR is similar to PDGFRTM, which gave us an indication that they might be monomeric as well, which would confirm results obtained for μOR in other studies [11, 12, 14].

### Single-molecule imaging does not show more dimers for κOR than for δOR and μOR

The split GFP approach indicated an increased dimerization tendency for κOR than for δOR and μOR, but an accurate quantification using this approach is impossible due to the stickiness of the GFP fragments. Also, the efficiency of GFP fluorescence recovery upon complementation is unknown, and therefore, the split GFP approach can only deliver a qualitative description. For these reasons, we chose a more direct approach and imaged green and red labeled receptors in the cell membrane on a single-molecule level. When the density of receptor molecules is sufficiently low to yield a large separation distance, then co-localization caused by random encounters in small, and co-localization of the fluorescent spots amounts to dimerization. Nevertheless, we will account for the impact of receptor density on the observed frequency of yellow spots appearing from random overlap. Since we abstained from the use of the split GFP and also introduced the mutation A206K that virtually eliminates the tendency of GFP to dimerize, the dimer fraction of the ORs should not be affected by interactions of the tags [20].

We fused a C-terminal monomeric GFP or an N-terminal SNAP-tag to the target proteins, hereby obtaining X-GFP and SNAP-X, where X was δOR, κOR, μOR, or PDGFRTM (the monomeric control). The SNAP-tag was labeled with a conjugate of the orange-red dye DY549-P1 and benzylguanine (SNAP549) (Fig. 2A). We recorded 488 nm / 561 nm excitation dual-color movies of CHO cells expressing one of the X-GFP/SNAP549-X pairs (Fig. 2B and Suppl. Movies 1, 2). We imaged cells with low surface densities of < 5 / µm^2^ of the proteins because with higher expression, individual molecules could not be separated from each other and random co-localization of the green and red emission from GFP and SNAP549 became too high. For all four protein pairs, we observed red and green fluorescent spots, and in a minor fraction of spots, red and green signals overlapped. The mobility of the spots was in the range described for ORs and other GPCRs in previous studies, and only a small fraction was immobile (Suppl. Note 2).

**Figure 2:**
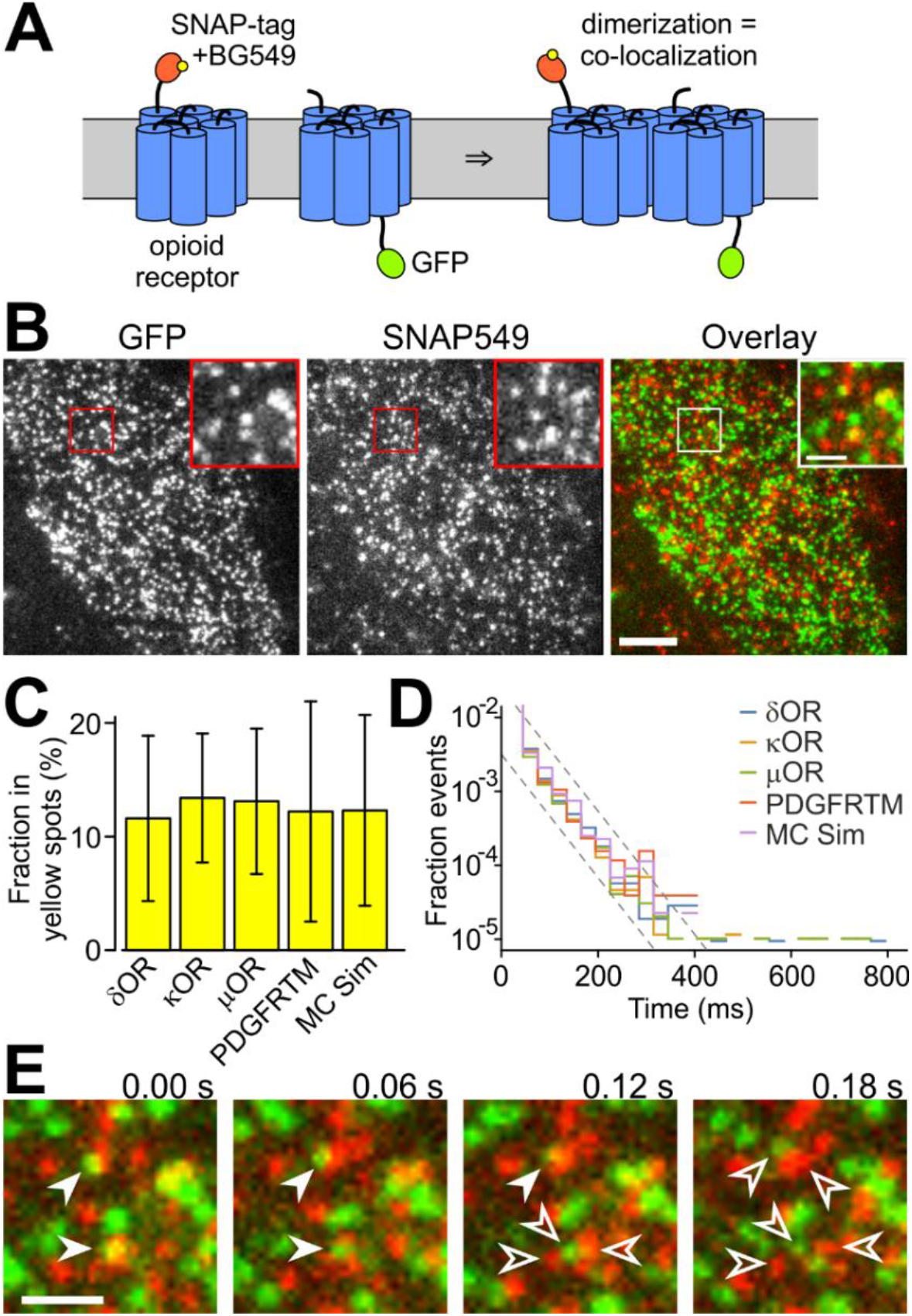
OR co-localization imaged on a single molecule level. **(A)** One receptor is labeled with the SNAP-tag and BG549, the other with GFP. When the receptors dimerize, the green and orange-red fluorescence co-localize. **(B)** Still image from a dual color movie. Inset shows magnified area. Scale bar 5 µm (inset 1 µm). **(C)** The fraction in yellow spots is similarly low for all ORs, the monomeric control, and the MC simulation. **(D)** The histogram of co-localization times decays with a time constant of about 30 ms for all proteins and the MC simulation. **(E)** Green and red spots that overlaid to yellow (filled arrowheads) moved apart within a few frames due to diffusion (empty arrowheads). Scale bars 1 µm.

To assess the overlap, we selected a representative area of the cell, where no major bright or empty areas were present, and tracked the number and positions of the spots in the green and red channels using an automated tracking algorithm [21]. We restricted the analysis to the first two fully illuminated frames, because photobleaching reduced the co-localization in later frames. Spots where the position in the two channels differed by 213 nm or less were classified as overlapping and referred to as ‘yellow’ in the following. The cutoff distance of 213 nm was determined from *bona fide* yellow spots in a positive control carrying both green and red tags (Suppl. Note 3). The average fraction of molecules in yellow spots was 12 ± 7% (s.d., n = 16 cells) for δOR, 13 ± 6% for κOR (n = 16), 13 ± 6% (n = 15) for μOR and 12 ± 10% (n = 6) for the negative control PDGFRTM (Fig. 2C). We observed a linear dependence of the yellow spot fraction on the receptor density, which supports the view that they were indeed a result of coincidental vicinity of non-interacting green and red spots (Suppl. Fig. 2A). Importantly, the parameters of linear fits of the yellow spot fraction to the spot density were not significantly different for the three ORs and the negative control and matched the analytical prediction of non-interacting proteins. Also, a Monte Carlo (MC) simulation of non-interacting spots yielded a similar fraction of yellow spots (12 ± 8%, n = 50).

We next evaluated the time that green and red spots remained co-localized in consecutive frames of the movies (excluding events with only a single frame of co-localization). For δOR, we obtained 98 ± 74 ms (s.d., n = 763 spots), for κOR 96 ± 85 ms (n = 584), for μOR 105 ± 94 ms (n = 581), and for PDGFRTM 102 ± 63 ms (n = 178). To further investigate possible mechanisms for the co-localization, we established a histogram of co-localization times. We used a semi-logarithmic presentation, where the rate constant of a decay process is visible as the histogram’s slope (Fig. 2D). After normalization to the total number of green and red localizations, the histograms for the three ORs, the PDGFRTM and the MC simulation virtually coincided, and more importantly, had a constant slope over most of their range. This suggests that a single process is responsible for co-localization and the loss of co-localization, which is the coincidental approach of green and red labeled receptors and their drifting apart due to diffusion (Fig. 2E). The time constant for loss of co-localization can be calculated from the estimate of the slope and is approximately 30 ms.

Although the time for co-localization was low for all four protein pairs, we sometimes observed yellow spots for all three ORs (but not for PDGFRTM) where green and red co-localized much longer (Suppl. Fig. 3A and Suppl. Movie 3). However, since they occurred very rarely, they had no impact on the average time of co-localization.

**Figure 3:**
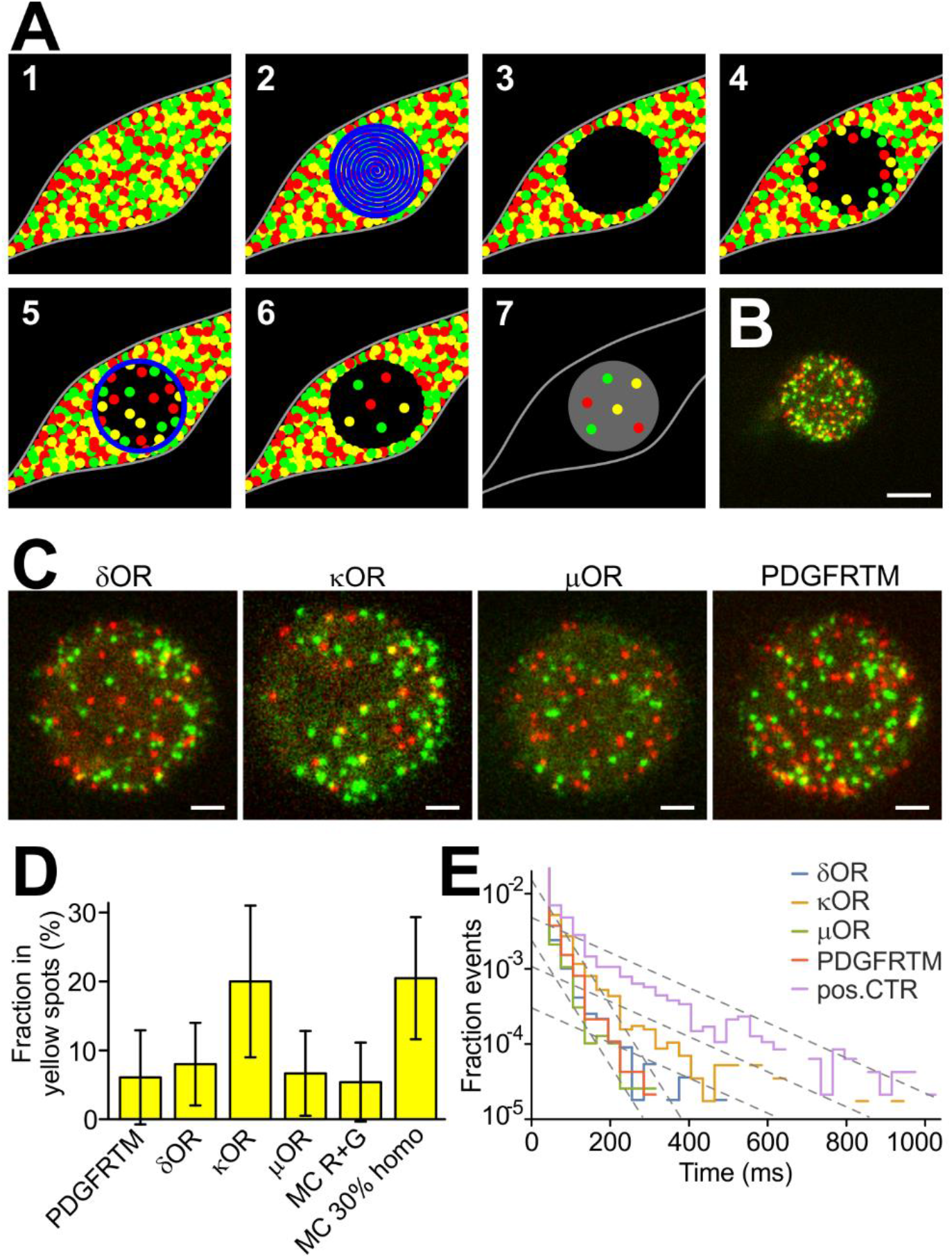
OR co-localization imaged with PhotoGate. **(A)** Schematic of the PhotoGate assay [15]. (1) A cell with a very high membrane density of fluorescent molecules (2) gets bleached in the central part, (3) leaving a dark bleached area. (4) After a while, the bleached area gets re-populated. (5) Bleaching a ring every few seconds prevents over-population by diffusion of unbleached molecules from the edges. (6) The result is a low density in the central area. (7) For imaging, only the central part gets illuminated. **(B)** First frame of a dual-color PhotoGate movie for κOR. Scale bar 5 µm. **(C)** Magnified central area for the ORs and PDGFRTM. For κOR, more yellow spots are visible. Scale bars 2 µm. **(D)** Fraction in yellow spots for the three ORs and the monomeric control PDGFRTM. **(E)** Time of co-localization of the green and red fluorescence in a yellow spot.

In the single-molecule tracking and co-localization, we did not observe a larger fraction of yellow spots, or a longer time of co-localization for the κOR than for the δOR, μOR, or the negative control PDGFRTM, in contrast to what we expected from the higher GFP-positive fraction in the split GFP assay. This suggests that under the low expression conditions used here, there was no significant level of dimerization for any of the proteins studied, and that the observed overlap was due to random co-localization without direct interaction.

### Single-molecule imaging of highly expressing cells with PhotoGate reveals κOR dimers

How can we resolve the apparent discrepancy between the results we obtained from the split GFP experiments, where κOR had a higher GFP-positive fraction than δOR and μOR, and the single-molecule imaging, where the fraction of yellow spots for κOR was similar to the other two ORs and the (presumably monomeric) control? One explanation that would reconcile these findings is that the dissociation constant for κOR is above the density from the single-molecule experiments, but below the density in the split GFP assay. Then, as we would expect for non-covalent binding, monomers and dimers (or higher-order clusters) can interconvert, and a higher density should shift the equilibrium to a higher dimer/cluster fraction.

To experimentally support this idea, we want to use a single-molecule co-localization approach that works at higher densities of molecules in the membrane. Two techniques called TOCCSL (Thinning Out Clusters while Conserving the Stoichiometry of Labeling) and PhotoGate were previously designed for this purpose [8, 16] (Fig. 3A). In a highly expressing cell, a part of the cell membrane gets photobleached quickly. After a recovery time, unbleached molecules re-populate the bleached area by lateral diffusion. The resulting density in the bleached area will be lower than the initial density. If the time allowed for recovery is shorter than the time for protein complexes to dissociate, the intact complexes can be imaged at lower density in the thinned-out region. In the PhotoGate technique, in addition to the initial bleaching exposure, a thin ring of high intensity is projected at certain time intervals to keep the density in the central imaging area low. In order to reduce glare from the highly expressing part of the cell, an iris is used to restrict the illumination used for imaging to the low expressing area.

We used the same constructs as for the previous experiment, i.e., we co-expressed X-GFP and SNAP549-X, where X was δOR, κOR, μOR, or PDGFRTM. However, this time we selected cells expressing a high density of molecules. In a small circular area of the cell with 10–15 µm diameter, GFP and SNAP549 were completely bleached with a focused laser beam. Then the laser was off for 5–20 s, depending on the expression level of the cell, to let molecules diffuse into the bleached area. Several rings were drawn at the outer edge of the bleached area to control the amount of recovery while allowing molecules inside the ring to distribute evenly. Finally, a two-color movie of the central area was recorded (Fig. 3B, C). Eventually, the remainder of the cell was imaged to determine the intensity per area and, by comparison to the unitary intensity of a single molecule, to calculate the density of molecules in the cell membrane.

As in the previous experiment, the overlap was assessed by automatically selecting green and red spots and classifying those spots as yellow where center positions were separated by the threshold distance of 213 nm or less. The average fraction of receptors in yellow spots was 8.0 ± 6.0% (s.d., n = 28 cells for δOR (avg. density in the unbleached cell 56 ± 37 / µm^2^), 20.0 ± 11.0% for κOR (n = 18, 53 ± 35 / µm^2^), 6.7 ± 6.1% (n = 23, 48 ± 35 / µm^2^) for μOR and 6.1 ± 6.8% (n = 20, 59 ± 32 / µm^2^) for PDGFRTM (Fig. 3D and Suppl. Movies 4–7). At high membrane densities, the κOR shows a significantly higher co-localization of green and red than δOR, μOR, or the PDGFRTM control (p < 0.00006). The dependence of the yellow spot fraction on density in the imaged region was similar for δOR, μOR, the monomeric control PDGFRTM and a MC simulation of non-interacting green and red spots (Fig. 3D and Suppl. Fig. 2B). However, the κOR showed a different behavior, which resembled a MC simulation of a homomeric protein where 35% of receptors resided in dimers and 65% in monomers.

To investigate the co-localization times of the ORs and the mechanisms involved, we analyzed the PhotoGate data in the same way as the initial single-molecule experiment without PhotoGate. We also imaged the construct SNAP549-PDGFRTM-GFP, where both tags are fused to the same protein (Fig. 3E). In the semi-exponential presentation, the histograms for δOR, μOR and the monomer control PDGFRTM (co-expressed SNAP549-PDGFRTM and PDGFRTM-GFP) again decay with a similar slope as for the experiment with low density, yielding a time constant for the loss of co-localization of 30 ms. The heterodimer mimic SNAP549-PDGFRTM-GFP also displays an initial decay with a similar slope, but then transitions into a slower decay of about 120 ms. For the κOR, we observe a similar behavior, with the slower decay starting a bit lower, but then continuing parallel to the decay of the heterodimer mimic. For SNAP549-PDGFRTM-GFP and κOR, we also observed multiple very long co-localizations (Suppl. Fig. 3B). The histograms for the heterodimer mimic and κOR can be interpreted as a superposition of two populations that have different mechanisms for the loss of co-localization. The faster process is the previously observed drifting apart due to diffusion. The other process, which is significantly slower, therefore is most likely the photobleaching or blinking of one of the tags, which causes the co-localization to end. The similarity of the slopes for the heterodimer mimic and κOR suggests that the mechanism is the same for both proteins.

The large fraction of yellow spots that shows a fast, diffusion-based loss of co-localization can be explained by co-localizations of partially photobleached complexes that appear in later frames of the 100-200 frame long movies. When we restricted the analysis to spots that were present in the first illuminated frame, the initial part of the histogram with the steeper slope nearly disappeared for the heterodimer mimic SNAP549-PDGFRTM-GFP, and became much smaller for κOR, while δOR and μOR still only showed the steep decay (Suppl. Fig. 3C).

We again observed a small fraction of immobile spots (around 5%, Suppl. Note 2), and wanted to test whether they contributed a major part to the yellow spots and the events with a longer lifetime. However, the yellow spots were distributed between mobile and immobile fraction in proportion to the fraction size, meaning that there was no correlation between yellow and immobile spots. After excluding the immobile spots, neither the fraction of yellow spots nor the lifetimes changed notably (Suppl. Fig. 4).

**Figure 4:**
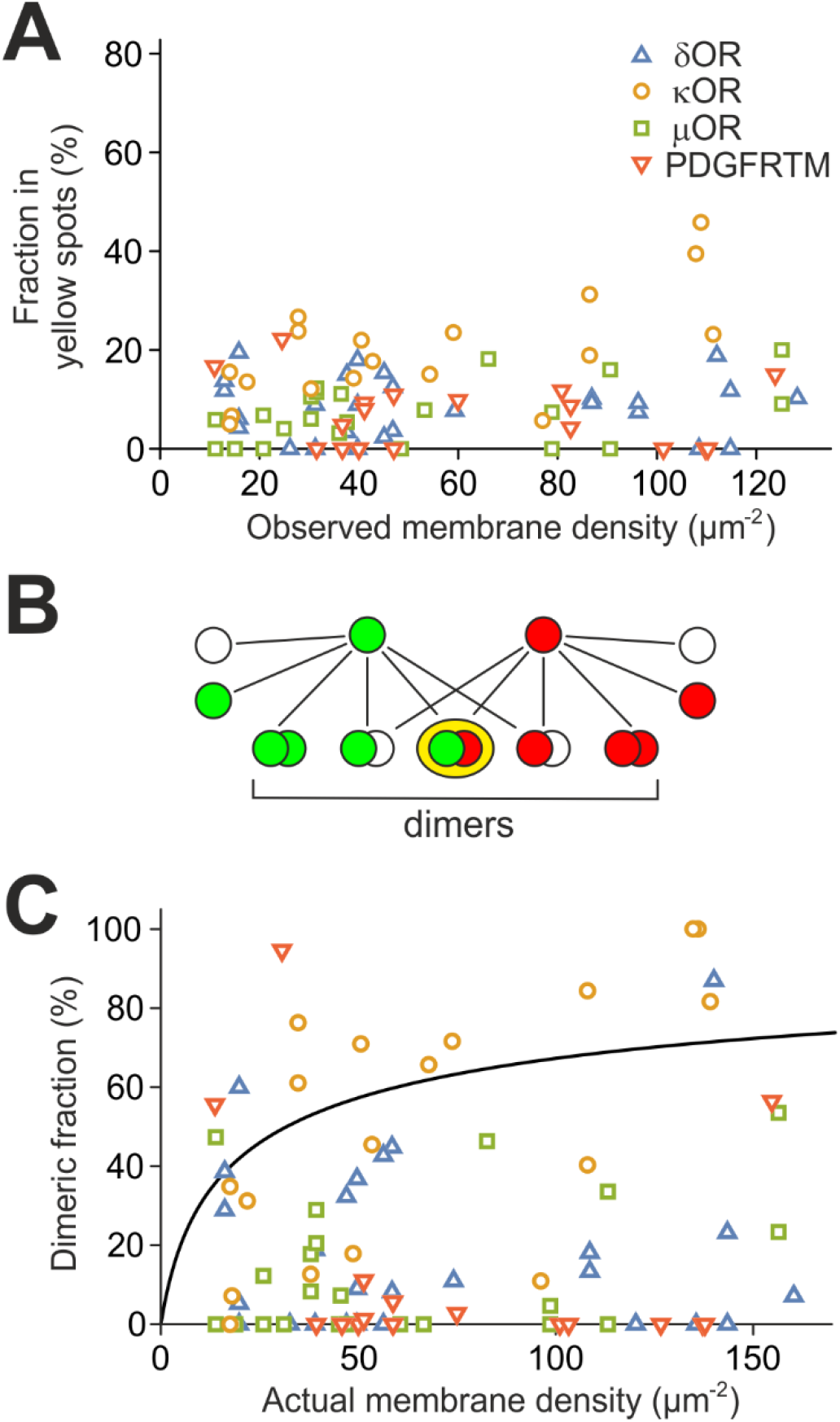
Dimerization of κOR. **(A)** The fraction of molecules in the yellow spots increases with density for κOR (green circles) but stays constant for δOR (red triangles), μOR (blue squares) and the monomeric control PDGFR (grey crosses). **(B)** The dimeric fraction is much larger than the fraction of molecules in yellow spots due to the non-fluorescent fraction of GFP, the unlabeled SNAP-tag, and green/green and red/red dimers. **(C)** The dimeric fraction of κOR increases up to 100%, but the dimeric fractions for δOR and μOR stay in the low range of the monomeric control. A fit of a binding curve (black line) to κOR yields the dissociation constant *k*_*d*_ = 32±15 / µm^2^.

These results support our assumption that an increased density shifts the equilibrium from monomers to dimers or higher-order clusters for κOR. Since PDGFRTM, δOR and μOR have no tendency to dimerize or cluster, they stay monomeric. Also, since there was a delay of 10-15 s between photobleaching and imaging, κOR dimers or clusters must be stable for at least 10 s, because with faster dissociation, we would not have observed any interactions.

### Dissociation constant of κOR

Finally, we set out to estimate the dissociation constant for κOR by analyzing how far the co-localization depended on the receptor density **before** reducing it by photobleaching. Therefore we plotted all values of the yellow fraction obtained from the PhotoGate experiments as a function of total spot density including green, red, and yellow before photobleaching (Fig. 4A). It is important to note that the density before photobleaching is not related to the spot density during imaging (which happens after photobleaching) and is up to 100x higher. We observe that for δOR, μOR and PDGFRTM, there is no trend for a higher yellow fraction at high densities. However, for κOR, the yellow fraction increases from a low value of 10% at a density of 14 / µm^2^ to a high value of 46% for densities of > 100 / µm^2^, supporting the expected density dependence of the κOR dimer/cluster fraction.

Importantly, the fraction of yellow spots did not exceed 50% at high densities, suggesting that no higher-order clusters were formed; in higher-order clusters, the chances are high to have both green and red labeled receptors, and therefore we would have expected the fraction of yellow spots to further increase and reach levels above 50%; in contrast, in homodimers, a maximum of 50% of the complexes can carry both colors because 25% should be green/green dimers and 25% red/red dimers. Therefore, we assume in the following that the κOR complexes are dimers.

The value shown in Fig. 4A is not the dimer fraction, but the fraction of yellow spots, which is not the same. While the fraction of yellow spots is limited due to green/green and red/red dimers and complexes containing non-fluorescent tags, we would expect the dimer fraction to reach up to 100%. To calculate the dimer fraction from the fraction of yellow spots, we established a model that accounts for random co-localization, monomer and dimer fractions, non-fluorescent GFP and unlabeled SNAP-tag, and the green/green and red/red dimers (Fig. 4B and Suppl. Note 4). We measured the fraction of non-fluorescent GFP and the fraction of unlabeled SNAP-tag in an experiment with SNAP549-PDGFRTM-GFP. Since every molecule carries both tags, the non-fluorescent/unlabeled fractions can directly be determined from the fraction of green, red, and yellow spots (Suppl. Note 5). The fraction of fluorescent GFP was 54 ± 3%, the fraction of labeled SNAP-tag was 62 ± 2% (n = 14, s.e.m.).

The model allowed us to calculate the fraction of receptor subunits in dimers from the numbers of yellow, green, and red spots, and the density during imaging (Fig. 4C). We find that for the κOR, the dimer fractions increase up to 100%, while for δOR and μOR, they remain on a lower level similar to PDGFRTM. This behavior becomes more visible when reducing variability by averaging several data points (Suppl. Fig. 5). We then set out to find a common dissociation constant that allows a best fit of the model parameters to the experimentally obtained data. The dissociation constant is an implicit parameter in the model, and is related to the fraction of receptors in dimers *f*_*d*_ by the equation

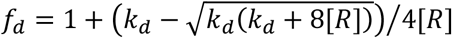

with the dissociation constant *k*_*d*_ and the receptor density [*R*], which we corrected for non-fluorescent GFP and unlabeled SNAP-tag (Suppl. Note 6). For the κOR, a nonlinear fit yielded the dissociation constant *k*_*d*_ = 32±15 / µm^2^ (68% confidence interval) (Fig. 4C). Similar attempts to determine dissociation constants for the other proteins yielded 8.4·10^3^ / µm^2^ for δOR, 9.4·10^3^ / µm^2^ for μOR and ∞ for PDGFRTM. Taken together, we conclude that κOR forms dimers at expression levels above 20 / µm^2^, but δOR and μOR stay monomeric, at least at densities up to 100 / µm^2^.

## Discussion

In this study we used a split GFP complementation assay, conventional single-molecule imaging, and the PhotoGate single-molecule technique to assess the dimerization of ORs. We find that κOR forms homodimers at densities below 100 / µm^2^, while δOR and μOR stay monomeric. The dissociation constant for κOR is *k*_*d*_ = 32±15 / µm^2^. We observe that co-localization of green and red labeled receptors of the δOR, the μOR and the monomeric control is only transient, whereas κOR forms longer lasting dimers at the same densities. Moreover, the lack of higher-order aggregates for high receptor densities suggests that the dimerization is a specific effect and not caused by coincidental co-localization. We also considered the possibility that co-localization was caused by clustering at cellular structures, e.g., clathrin coated pits; in this case, we would expect immobilization of the spots. However, the immobile fraction of spots was smaller for the experiment with PhotoGate than for the experiment without PhotoGate, suggesting that the co-localization was not caused by clustering at cytoskeletal structures or internalization sites (Suppl. Note 2).

In general, it is difficult to measure lateral affinities of membrane proteins because of the inability of biochemical methods to determine the density of the target protein in the plasma membrane. So far, this can only be achieved with fluorescence-based assays where the signal from the labeled protein in the membrane of a living cell can be compared to a reference of known concentration or a direct observation of single molecules is possible. With conventional single-molecule imaging, maximum densities of 5 / µm^2^, are accessible [9,10]. With bulk fluorescence methods using confocal microscopy, the minimum measured densities were 160 / µm^2^ [22–24]. The PhotoGate approach we used covers the missing range of 10–100 / µm^2^. It remains to be seen if measurements from the three ranges can be consolidated into a common framework.

A major contribution to the error in the dissociation constant originates – as for many single-molecule methods – from the uncertainty inherent to counting low numbers of events, in our case the green, red, and yellow spots. The uncertainty in the initial counts (Fig. 4A) gets even amplified when subtracting the expected number of yellow spots caused by random co-localization (Fig. 4C). A second error source that is difficult to account for is the variability of the SNAP-tag labeling efficiency. In the model to calculate the fraction of dimers from the fraction of yellow spots, we used the average SNAP-tag labeling efficiency, but the actual labeling efficiency might vary from one cell to another, e.g., due to different accessibility of the substrate in the extracellular solution to the cell.

A possible source for a systematic error that is more difficult to compensate lies in the design of the PhotoGate technique. A delay is required between the initial bleaching and the movie acquisition to allow molecules from the outside to diffuse into the bleached center (Fig. 3A). In our experiments, this delay was in the range of 10–20 s. If receptor dimers dissociate in a comparable time frame, then a significant fraction of the dimers will have decayed by the time of imaging and will therefore appear as monomers. However, we estimate that this effect reduces the dimer fraction by less than 10% in the case of the κOR since we obtained a large dimer fraction for high membrane densities. On the other hand, the fact that even after a long delay of up to 20 s κOR dimers remain intact, means that the κOR dissociation time is larger or at least in the range of 20 s. In principle, it is possible that the δOR and/or μOR dimer fractions are affected by fast dissociation into monomers (at least faster than 10 s), and that our interpretation of δOR and μOR to be monomers at all tested densities is incorrect. However, one would then expect the GFP positive fraction in the split GFP assay to be larger than what we observed. Therefore, we think that our experiments are consistent with δOR and μOR being monomers at the densities we observed.

For many other *in vitro* studies, the receptors were expressed under control of the strong CMV promoter; in contrast, we here used a promoter with inducible expression and selected cells with receptor densities of 200 / µm^2^ or less. To investigate whether the different expression levels could be a possible cause for the apparent discrepancies, we measured the membrane densities of δOR when expressed under the CMV promoter. We found that although the majority of cells expressed less than 500 receptors/µm^2^, receptor densities reached up to 6000 / µm^2^ (Suppl. Fig. 6). But because the higher expressing cells contain more receptors, only a small fraction of 4% of the receptors experience densities below 500 receptors/µm^2^, and more than half of the receptors reside at densities above 3000 / µm^2^. Accordingly, bulk experiments (e.g., BRET and co-IP) will mainly yield the signals from the small fraction of high-density cells. Our observation that δOR and μOR are monomeric could be therefore reconciled with other studies if the dissociation constants of δOR and μOR lies above 200 / µm^2^.

Some current sources offer data on RNA levels in all organs, which suggest that ORs are expressed in most tissues, but at strongly differing levels [25]. However, it was not determined whether just a small fraction of cells express the receptors strongly, or most of the cells at low or moderate levels. Accordingly, expression levels and membrane densities in individual cells remain unknown. To understand the impact of OR dimerization *in vivo*, receptor membrane densities in different cell types in the body need to be measured in the future.

## Materials and Methods

### DNA constructs

The murine δ-and κ-opioid receptor, rat µ-opioid receptor, and the transmembrane domain of PDGFRα were cloned into the pWHE636 vector (gift by Christian Berens) that contains a tetracycline inducible promoter [26]. An N-terminal signal peptide and the SNAP-tag were fused to the N termini and GFP to the C-termini of the receptors and PDGFRTM with flexible linkers. In order to have well-folding, soluble domains on both sides of PDGFRTM, we added a non-fluorescent mEGFP carrying the mutation Y66l (to render it non-fluorescent) to the C-terminus of SNAP-PDGFRTM, and a signal peptide, followed by a HALO-tag, to the N-terminus of PDGFRTM-GFP. For split GFP, β-sheets 1–10 (aa 1–214) of a GFP variant were fused to the constructs with the SNAP-tag, and β-sheet 11 (aa 215–230) and mKate to the constructs without SNAP-tag [19].

### Expression of ORs in CHO cells

For inducible expression in mammalian cells, a stable CHO-K1 cell line (ATCC) was made with the regulator plasmid pWHE644 containing a tetracycline-regulated transactivator and a transcriptional silencer [26]. The cells were seeded on a high refractive index coverslip (n = 1.78) and grown in complete DMEM for 12–15 h to obtain 60–80% confluency before transfection with polyethylenimine (1 mg/ml) with 1 µg of each DNA per coverslip. After 3 h, cells were washed in DPBS and incubated with doxycycline in DMEM to induce expression. The induction was terminated by washing with DPBS, and then complete DMEM was added to cells. Imaging started after 12–15 h of induction of expression. For single-molecule imaging and split-GFP assay, cells were incubated for 2 h with 1 µg/ml of doxycycline hyclate, for the PhotoGate experiment for 3 h with 2 µg/ml. For SNAP-tag labeling, cells were incubated with 2 nM benzylguanine-Alexa Fluor 647 (New England Biolabs, SNAP-Surface Alexa Fluor 647) or custom synthesized SNAP substrate benzylguanine-DY549P1 for 30 min before an experiment, and unbound dye was washed away by rinsing 5x with DPBS.

### Dye synthesis

HPLC analysis (1 mL/min) and purification (3 mL/min) were performed on an Agilent Technologies 1260 Infinity system using UV detection at 290 nm and – for analysis – a Phenomenex Kinetex® 5u XB-C18 100 Å 250 × 4.6 mm column or – for purification – a Phenomenex Synergi® 10u Hydro–RP 80 Å 250 × 15.0 mm column. Eluent A was water containing 0.05 % trifluoroacetic acid (TFA) and eluent B was acetonitrile containing 0.05 % TFA. Linear gradient conditions were as follows: 0–1 min, A/B (90:10); 1−21 min, linear increase to 100 % of B; 21−23 min, 100 % B; 23– 23.3 min: A/B (90:10); 23.3–26 min: A/B (90:10). Characterization was performed through mass spectrometry and mass spectra were recorded on a Thermo Scientific Exactive mass spectrometer using electrospray ionization (ESI) as ion sources.

To a stirred solution of 6-((4-(aminomethyl)benzyl)oxy)-7*H*-purin-2-amine (1.4 mg, 5.2 µmol, 1.1 eq.) in CH_3_CN (250 µL), under inert atmosphere, was added the NHS ester of DY549-P1 (5 mg, 4.8 µmol, 1 eq.) in 250 µL of H_2_O, followed by NaHCO_3_ (50 µL of a 1 M solution: final concentration of c.a. 0.1 M) and the reaction was stirred protected from light for 48 h. After completion (RP-C18 TLC: H_2_O/CH_3_CN 3:7) the solvent was removed to give a pink oil. The crude SNAP549 was then dissolved in the minimum volume of methanol, and purified using preparative HPLC (see conditions above). The product was isolated as a pink oil (3.3 mg, 55 % yield). C_49_H_57_N_8_Na_3_O_15_S_4_ (1195.25 g/mol). HPLC: t_R_ = 6.935 min (82 % purity – 18 % of DY549-P1-OH remaining). ESI-HRMS(–): m/z calcd for C_49_H_56_N_8_Na_3_O_15_S_4_: 1193.2447 [M–H]^−^; found 1193.2434 [M–H]^−^; m/z calcd for C_49_H_57_N_8_Na_2_O_15_S_4_: 1171.2627 [M–Na]^−^; found 1171.2611 [M–Na]^−^; m/z calcd for C_44_H_54_N_3_Na_2_O_15_S_4_: 1038.2239 [M–2AP–Na]^−^; found 1038.2229 [M–2AP–Na]^−^.

### Microscopy

Imaging was done on an Olympus IX71 base equipped with an Olympus 100x NA1.70 objective, a back-illuminated EMCCD camera (Andor iXon DV-897 BV), an emission filter wheel and an OptoSplit II beam splitter (Cairn). GFP, mKate/BG549, and BG647 were excited through a custom built total internal reflection illumination pathway either consecutively (split GFP assay) or in alternating excitation (single-molecule movies). Movies (256×256 pixels) were recorded at a frame rate of 17 or 33 Hz or, for alternating excitation, at a frame rate of 65 Hz. Single molecule imaging was done at power densities of 100–250 W/cm^2^. During the experiments, cells with similar expression levels in all channels were chosen. For single-molecule imaging without PhotoGate, cells with expression < 5 / µm^2^ were selected.

### PhotoGate

The PhotoGate assay was done similarly as described in [16]. A 473 nm laser beam (25 mW before the objective) focused on the plane of the plasma membrane was directed using a galvo scanning system placed at an appropriate position in the light path. Despite 473nm being far from the excitation maximum for DY549-P1, the intensity of the laser was sufficiently high to photobleach the dye completely. A region of interest of diameter 10–15 µm was bleached for about 10 s, depositing a total energy of 2.5 mJ/µm^2^, by moving the focused beam in a spiral motion. After a 5–20 s delay (depending on fluorophore density), 2–8 rings were drawn on the cell’s surface at intervals of 3–5 s, to control the density inside the pre-bleached area. Right after, an iris in the illumination pathway was closed to restrict the illumination to the central area and eliminate glare from the bright area outside, and the movie acquisition was started.

### Analysis of single-molecule images

The first two fully illuminated frames for each color were analyzed. A rectangular region of interest was chosen that was completely covered by plasma membrane and had an even distribution of spots. The center positions of green and red spots were automatically selected by the MOSAIC tracking suite [21]. The fraction of receptors in dimers was calculated as *f*_*d*_ = 2*N*_*y*_/(2*N*_*y*_ + *N*_*g*_ + *N*_*r*_). When the centers of green and red spots were as close as 213 nm or less, both were combined to form a yellow spot. Significance levels were calculated using the Mann-Whitney U test. For determination of co-localization time constants, green and red trajectories identified by the MOSAIC tracking plugin were overlaid. The “yellow” trajectory was interrupted as soon as the difference in the positions of the green and red spot was larger than 213 nm or one of the trajectories ended.

### Density evaluation

For calculating the density of fluorescent receptors in the split-GFP and the PhotoGate assay, the intensities of single molecules were compared to the intensities of a region of interest. First, a small region of interest (0.5×0.5 µm^2^) around individual fluorescent molecules (either from a lower-expressing cell or after photobleaching of the majority of spots) was selected, and background from a nearby region without a spot or the same region after photobleaching of the spot was subtracted. An average value was formed from 10–20 spots. Similarly, for an area with high density, background was selected from an area outside the cell.

## Author Contributions

K. C., and M. U. conceived the study, designed experiments, and wrote the manuscript. C. L., K. C., M. M. and M. U. established the analysis of the data. N. P. F. B. and M. J. established the synthesis and synthesized the fluorescent SNAP-tag substrate.

## Acknowledgements

This work was supported by Deutsche Forschungsgemeinschaft (DFG) grants UL 312/6-1, RTG 2202, and CRC 992, the Excellence Initiative of the German Federal and State Governments (EXC 294), and under Germany’s Excellence Strategy (EXC-2189 – Project ID: 390939984), the Grant Agency of the Czech Republic (GA17-05903S) and the Faculty of Science, Charles University in Prague (SVV260427/2020, FM/a/2017-2-072, SVV260427/2018).

