## Supplementary Material for "Kappa but not delta or mu opioid receptors form homodimers at low membrane densities"

Supplementary Figure 1 3
Single-molecule events in the split GFP assay

Supplementary Figure 2 4
Density dependence of the yellow spots fraction

Supplementary Figure 3 5
Long co-localization events for δOR, κOR, μOR

Supplementary Figure 4 6
Yellow spots fraction and lifetimes after excluding immobile spots

Supplementary Figure 5 7
Dimer fraction after averaging

Supplementary Figure 6 8
Receptor membrane densities from expression under the CMV promoter

Supplementary Note 1 9
Calibration of GFP reconstitution in the split GFP assay

Supplementary Note 2 12
Fraction of immobile spots and diffusion coefficient

Supplementary Note 3 14
Determination of threshold distance for green and red to be considered co-localized

Supplementary Note 4 15
Model to calculate dimer fraction from yellow fraction

Supplementary Note 5 17
Model to calculate fluorescent GFP and labeled SNAP-tag fractions

Supplementary Note 6 18
Fraction of receptor subunits in dimers in dependence on total subunit density

Supplementary Note 7 19
Fitting of dissociation curve to κOR yellow spot fraction

Supplementary Movie 1
Cell expressing μOR-GFP and SNAP549-μOR (scale bar 5 μm)

Supplementary Movie 2
Cell expressing μOR-GFP and SNAP549-μOR (close-up, scale bar 1 μm)

Supplementary Movie 3
Cell expressing κOR-GFP and SNAP549-κOR (close-up, scale bar 1 μm)

Supplementary Movie 4
Cell expressing δOR-GFP and SNAP549-δOR (PhotoGate, scale bar 2 μm)

Supplementary Movie 5
Cell expressing κOR-GFP and SNAP549-κOR (PhotoGate, scale bar 2 μm)

Supplementary Movie 6
Cell expressing μOR-GFP and SNAP549-μOR (PhotoGate, scale bar 2 μm)

Supplementary Movie 7
Cell expressing PDGFRTM-GFP and SNAP549-PDGFRTM (PhotoGate, scale bar 2 μm)

#### Supplementary Figure 1

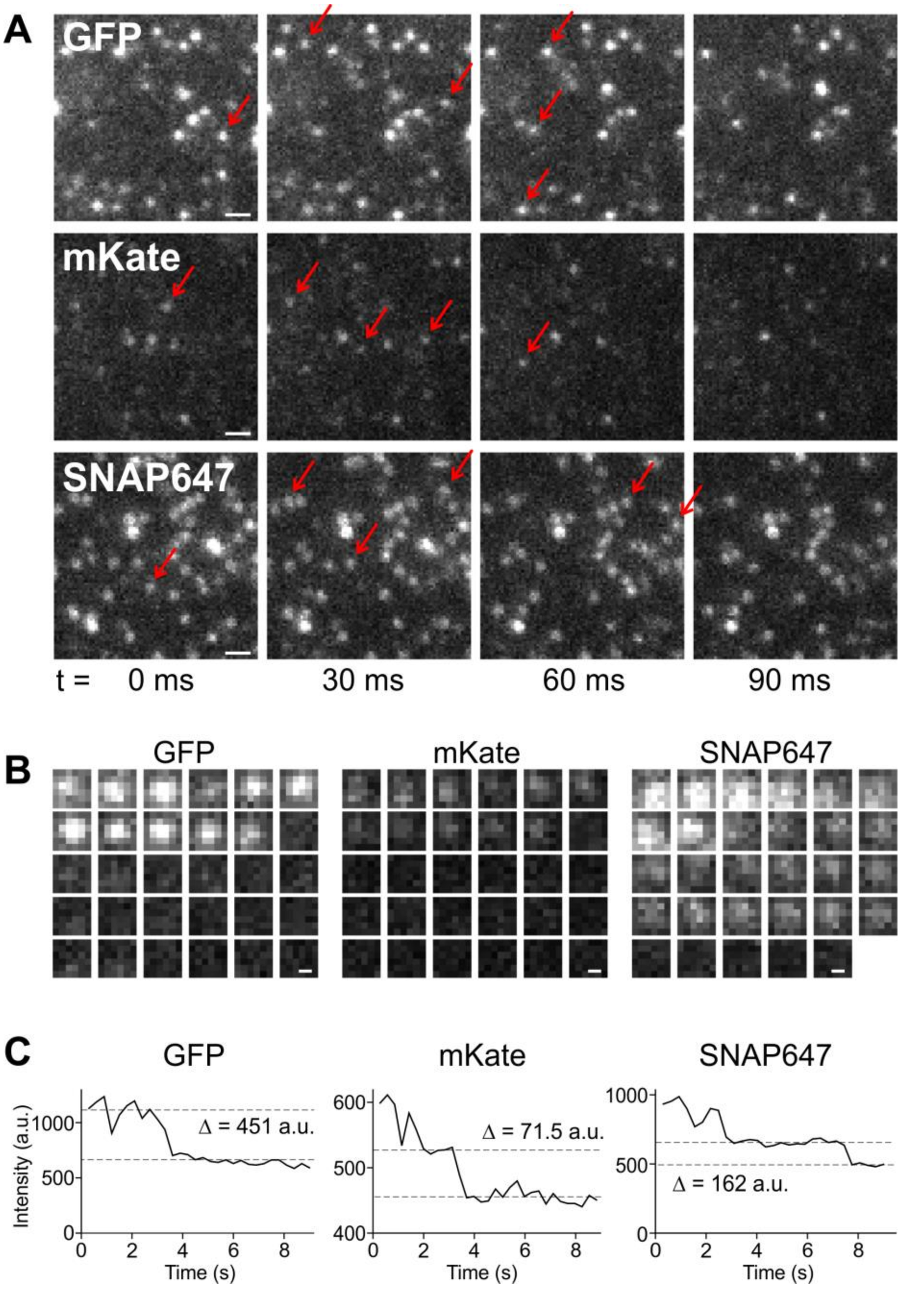


**Supplementary Figure 1: Single-molecule events in the split GFP assay. (A)** Magnified view of a CHO cell expressing low densities of SNAP-KOR-GFP10 and KOR-GFP11-mKate: In the four consecutive frames, red arrows mark spots that photobleach are not visible in the next frame any more. Scale bar 1 µm. **(B)** Time course of representative individual spots in consecutive 30 ms frames. Scale bar 200 nm. **(C)** The intensity time course displays distinct drops, suggesting that the spots are single molecules. The intensity levels before and after the last photobleaching event are marked by dashed lines.

#### Supplementary Figure 2

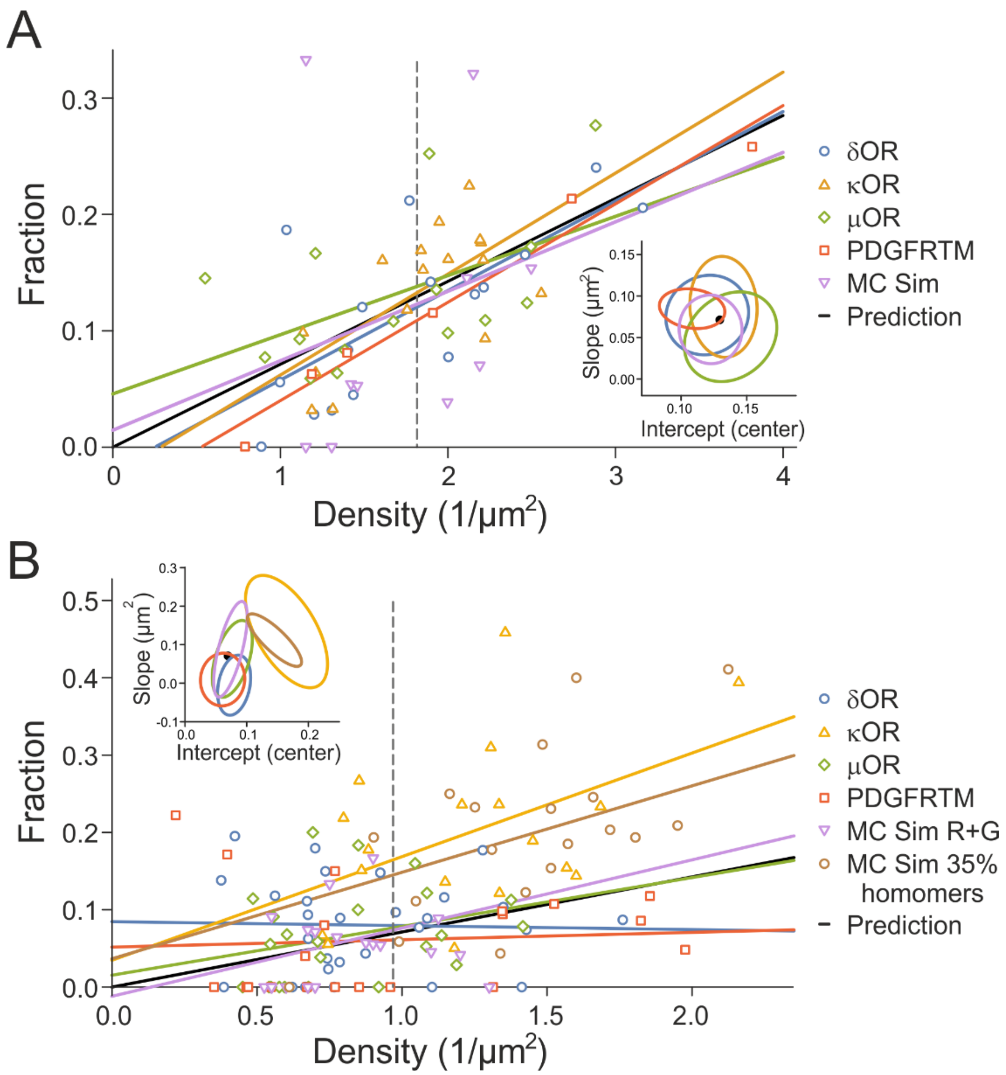


**Supplementary Figure 2: Density dependence of the yellow spots fraction. (A)** For single-molecule imaging at low density. The fraction of yellow spots increases with the receptor density. The linear fits (solid lines) of the ORs almost coincide with the fits for the negative control PDGFRTM and data from a Monte Carlo simulation. The dashed line indicates the average density of the four experimental distributions. The inset shows the 90% confidence regions of the two fit parameters: the slope of the fit line and the intercept value with the dashed line in the center of the experimental distributions. The black line and dot indicate the analytically derived estimate for the yellow fraction $f(d) = d/2 \cdot \pi \cdot r_0{}^2$, where $d$ is the density and $r_0 = 213\ \mathrm{nm}$ is the distance for a green and red spot to be considered co-localized. The overlap of the confidence regions and the analytic prediction supports the view that for all proteins, the fraction of yellow spots is caused by the same mechanism, namely coincidental overlap of green and red. **(B)** For single-molecule imaging after PhotoGate. For κOR, the density of yellow spots is increased and similar to a MC simulation of 35% dimers and 65% monomers.

#### Supplementary Figure 3

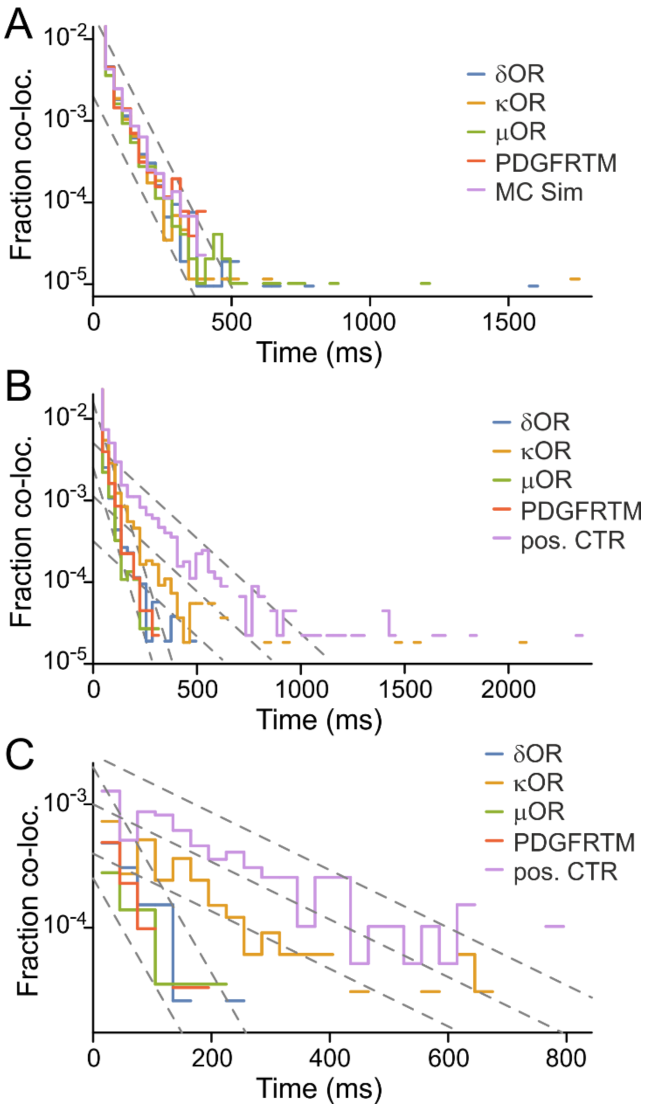


**Supplementary Figure 3: Co-localization times. (A)** Single-molecule experiment without PhotoGate. Very few long-lasting co-localizations exist for the ORs, but not for the PDGFRTM or in the MC Simulation. Since the probability for these events to happen from random co-localization and coincidental co-diffusion is very low, it is more likely that they are caused by an interaction of green and red labeled ORs. **(B)** Single-molecule experiment with PhotoGate. The heterodimer-mimicking construct SNAP549-PDGFRTM-GFP and κOR, but not δOR, μOR or the monomer control PDGFRTM (co-expressed SNAP549-PDGFRTM and PDGFRTM-GFP) show some events with long-lasting co-localization. **(C)** Co-localization events that started in the first illuminated frame. Guidelines for all panels have slopes of 30 ms and 120 ms.

#### Supplementary Figure 4

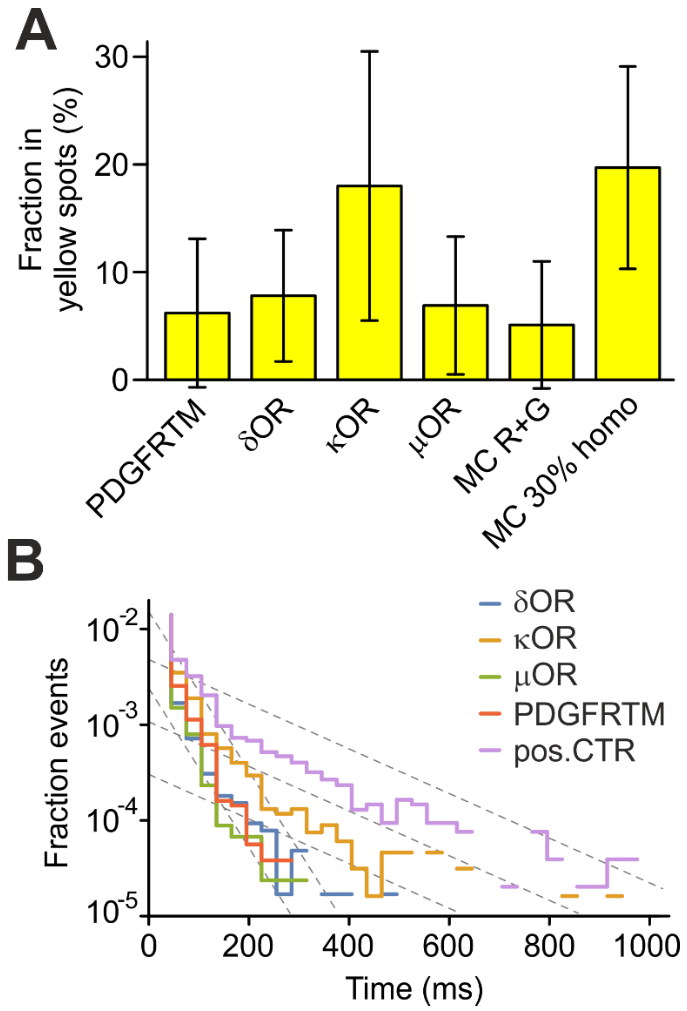


**Supplementary Figure 4: Yellow spots fraction and lifetimes after excluding immobile spots.** After exclusion of the immobile spots, **(A)** the fraction in yellow spots and **(B)** the lifetimes for the PhotoGate experiments did not change notably.

#### Supplementary Figure 5

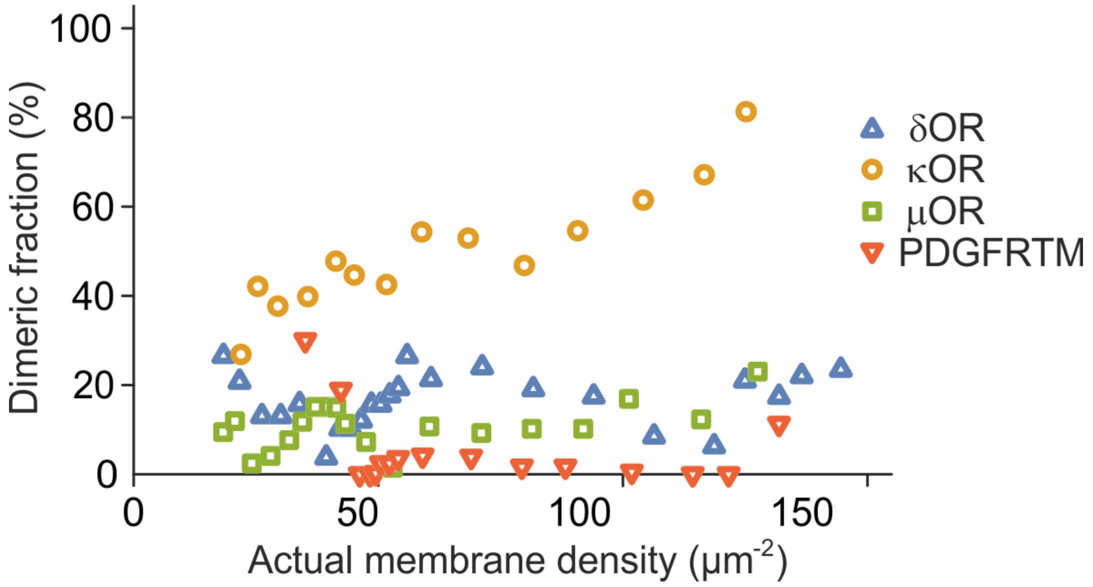


**Supplementary Figure 5: Dimer fraction after averaging.** When 5 consecutive data points are averaged, the trend of κOR to increase at higher densities, and the low dimer fractions for δOR, μOR and PDGFRTM become more visible.

#### Supplementary Figure 6

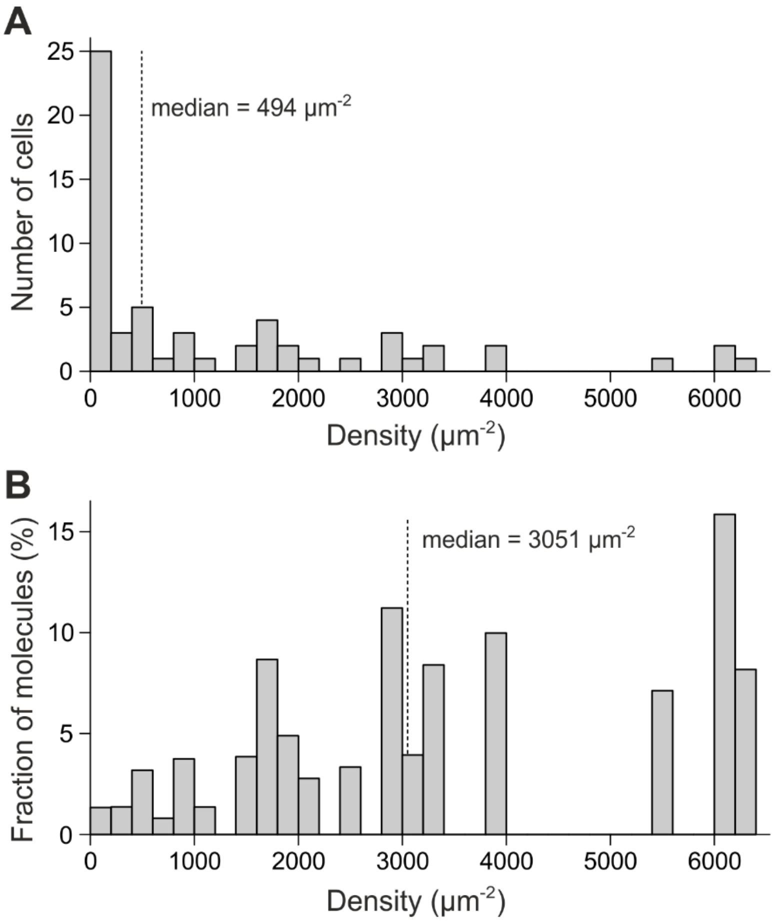


**Supplementary Figure 6: Receptor membrane densities from expression under the CMV promoter. (A)** Histogram of δOR-GFP receptor densities at the plasma membrane taken from 60 cells. 50% of the cells express at densities below 500 receptors/μm$^2$, and only 15% of the cells express at 3000 receptors/μm$^2$ or more. **(B)** A histogram of all receptors expressed at different densities, from the same cells as in (A). Only 4% of the receptors are at membrane densities below 500 receptors/μm$^2$, and 53% of the receptors reside at 3000 receptors/μm$^2$ or more.

#### Supplementary Note 1

##### Calibration of GFP reconstitution in the split GFP assay

We calculated the GFP-positive fraction as the ratio of GFP density to the smaller value of mKate and SNAP647 densities. The rationale behind this approximation is given in the following. Hereby, mKate-labeled proteins stand for X-GFP11-mKate, and SNAP-labeled proteins stand for SNAP-X-GFP10.

Our assumption is that the protein of interest, although primarily dimeric, is in equilibrium between monomers and dimers. Even when the equilibrium is strongly shifted towards dimers at high densities, we assume that the molecules associate and dissociate multiple times between protein synthesis and the experiment, if possible (once fluorescence has been reconstituted, dissociation is not possible anymore). A higher number of consecutive association/dissociation cycles will lead to a higher degree of reconstitution, because molecules that were part of mKate-mKate or SNAP-SNAP dimers (where reconstitution is not possible), have additional chances to be incorporated into mKate-SNAP dimers (where reconstitution will occur).

Examples for 50% mKate- and 80% mKate-labeled protein (the remaining part is SNAP-labeled) and the first three association/dissociation cycles show that after a few cycles, the majority of the lower-expressing species is incorporated into the stable dimers, which are covalently linked due to the GFP reconstitution (Suppl. Fig. 6A, B). Therefore, it appears appropriate to use the smaller value of mKate and SNAP647 densities as the reference to determine the degree of GFP reconstitution.

In contrast, when we would use e.g. the average of the mKate and SNAP647 densities, an error would be introduced, because a certain fraction of the higher expressing species remains without a partner for reconstitution. The error introduced increases with increasingly different expression levels of the mKate- to the SNAP-labeled species (Suppl. Fig. 6C).

We calculated the degree of GFP reconstitution after n association/dissociation cycles with the following model. The initial fraction of free molecules is $f_{fr,1} = 1$, and the initial fraction of mKate-labeled protein is $f_{K,1} = N_K/(N_K + N_S)$, where $N_K$ and $N_S$ are the numbers of mKate-labeled and SNAP-labeled molecules. After the first association cycle, the fractions for mKate-mKate dimers $f_{KK,1}$, SNAP-SNAP dimers $f_{SS,1}$ and mKate-SNAP dimers $f_{KS,1}$ are

$$\begin{aligned} f_{KK,1} &= f_{fr,1} \cdot f_{K,1} \cdot f_{K,1} \\ f_{SS,1} &= f_{fr,1} \cdot (1 - f_{K,1}) \cdot (1 - f_{K,1}) \\ f_{KS,1} &= f_{fr,1} \cdot 2 \cdot f_{K,1} \cdot (1 - f_{K,1}) \end{aligned}$$

Since the mKate-SNAP dimers are covalently bound to each other after the association, they will not dissociate again, but the mKate-mKate and SNAP-SNAP dimers can dissociate to monomers. Accordingly, the fraction of free molecules is $f_{fr,2}$ and within these, the fraction of mKate-labeled protein $f_{K,2}$ are:

$$\begin{aligned} f_{fr,2} &= f_{KK,1} + f_{SS,1} \\ f_{K,2} &= \frac{f_{KK,1}}{f_{KK,1} + f_{SS,1}} \end{aligned}$$

The fractions of mKate-mKate dimers, SNAP-SNAP dimers and mKate-SNAP dimers in the following association cycles can be obtained from an iterative application of these formulas:

$$f_{KK,i} = f_{fr,i} \cdot f_{K,i} \cdot f_{K,i}$$
$$f_{SS,i} = f_{fr,i} \cdot (1 - f_{K,i}) \cdot (1 - f_{K,i})$$
$$f_{KS,i} = f_{fr,i} \cdot 2 \cdot f_{K,i} \cdot (1 - f_{K,i})$$

$$f_{fr,i+1} = f_{KK,i} + f_{SS,i}$$
$$f_{K,i+1} = \frac{f_{KK,i}}{f_{KK,i} + f_{SS,i}}$$

The total number of molecules is $N_K + N_S$, and, since two molecules form one GFP, the number of green molecules after n cycles of association and dissociation is

$$N_{G,n} = \frac{N_K + N_S}{2} \cdot \sum_{i=1}^{n} f_{KS,i}$$

The ratio of GFP molecules after n cycles of association and dissociation to the smaller value of mKate and SNAP647 molecules is:

$$f_{min}(n) = N_{G,n}/\mathrm{Min}(N_K, N_S)$$

whereas the ratio of GFP molecules after n cycles of association and dissociation to the average of mKate and SNAP647 molecules is

$$f_{avg}(n) = N_{G,n}/\mathrm{Avg}(N_K, N_S)$$

While $f_{avg}(n)$ is becoming small when expression levels of mKate- and SNAP647-labeled molecules are very uneven, the function $f_{min}(n)$ is close to unity for all expression levels, especially when the number of association/dissociation cycles is large (Suppl. Fig. 6C). For a protein that is preferentially dimeric, but can cycle between monomers and dimers, our model therefore suggests that $f_{min}(n)$, i.e. the ratio of the GFP density to the smaller value of mKate and SNAP647 densities is well suited to assess the degree of GFP reconstitution.

For a monomeric protein, since there is (in theory) no association, $N_{G,n}$ is zero, and therefore, also $f_{min}(n)$ and $f_{avg}(n)$ both remain zero.

**Supplementary Figure 6 (next page): Degree of reconstitution in the split GFP assay. (A)** An example calculation for 50% mKate and 50% SNAP-labeled molecules for the first three association/dissociation cycles. After three cycles, 87.5% of mKate-labeled molecules are incorporated into the covalently fused dimers with reconstituted GFP. **(B)** An example calculation for 20% mKate and 80% SNAP-labeled molecules for the first three association/dissociation cycles. After three cycles, 99.995% of mKate-labeled molecules are incorporated into the covalently fused dimers with reconstituted GFP. **(C)** Model calculations demonstrate that for more than 2 association/dissociation cycles, the ratio of the GFP density to the smaller value of mKate and SNAP647 densities is well suited to assess the degree of GFP reconstitution. Only for equal expression of mKate and SNAP647, using the average of mKate and SNAP647 instead of the minimum would give a similar performance.

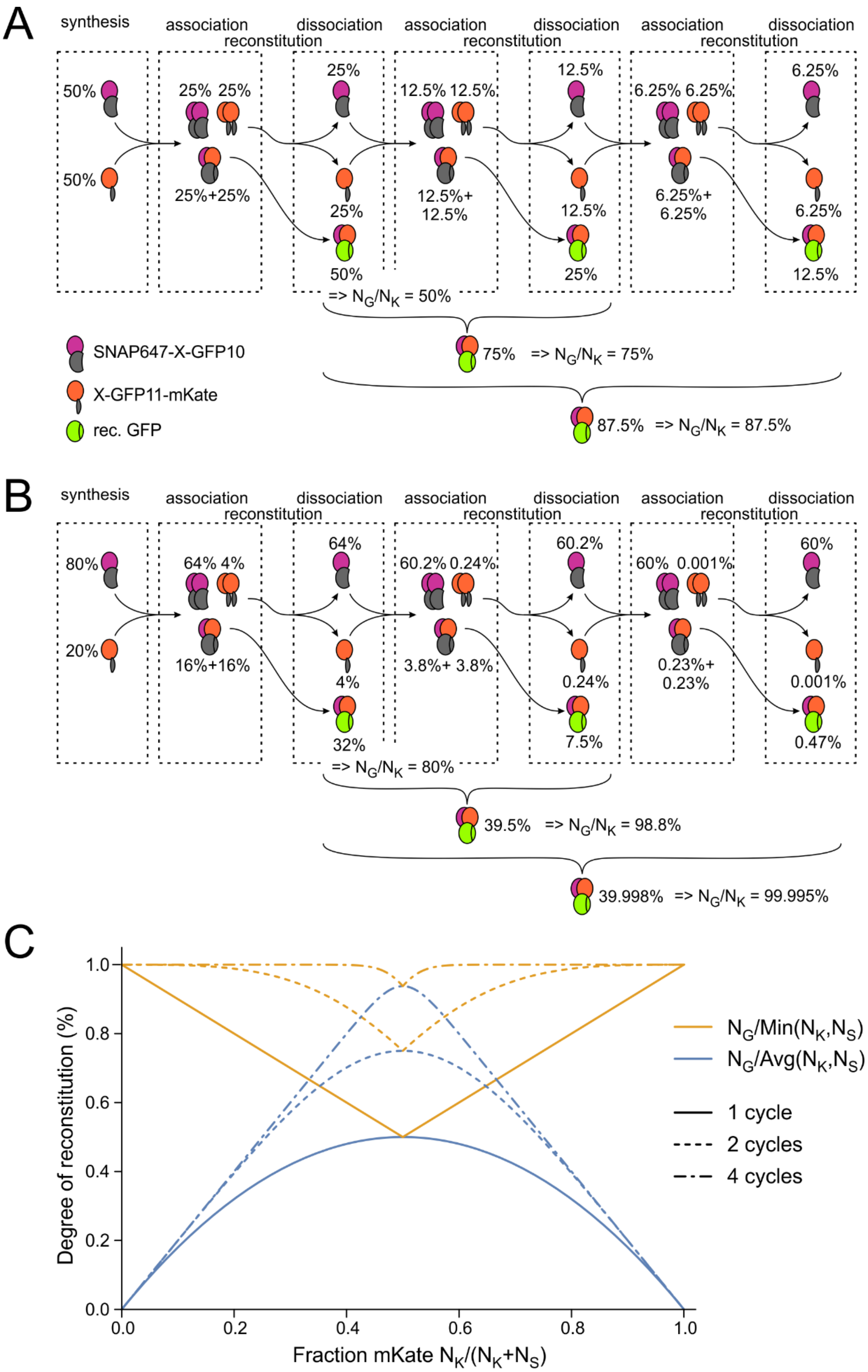
A
synthesis
association
reconstitution
dissociation
50%
25% 25%
25%+25%
25%
50%
=> $N_G/N_K$ = 50%
12.5% 12.5%
12.5%+
12.5%
6.25% 6.25%
6.25%+
6.25%
6.25%
12.5%
75% => $N_G/N_K$ = 75%
87.5% => $N_G/N_K$ = 87.5%
SNAP647-X-GFP10
X-GFP11-mKate
rec. GFP
B
80%
20%
64% 4%
16%+16%
64%
4%
32%
=> $N_G/N_K$ = 80%
60.2% 0.24%
3.8%+ 3.8%
60.2%
0.24%
7.5%
60% 0.001%
0.23%+
0.23%
60%
0.001%
0.47%
39.5% => $N_G/N_K$ = 98.8%
39.998% => $N_G/N_K$ = 99.995%
C
Degree of reconstitution (%)
Fraction mKate $N_K/(N_K+N_S)$
$N_G/Min(N_K,N_S)$
$N_G/Avg(N_K,N_S)$
1 cycle
2 cycles
4 cycles

#### Supplementary Note 2

##### Fraction of immobile spots

To first find a criterion for immobility, we selected 10 green and 10 red spots from single-molecule movies of the δOR, κOR, μOR and PDGFRTM constructs which we visually assessed to be immobile, with lengths between 14 and 121 frames (average 47 frames, frame time 30 ms). We found that their radius of apparent movement (due to localization inaccuracy) was 90 ± 20 nm with a maximum of 123 nm. Therefore, we used 125 nm as a fixed cutoff criterion for immobility, such that the localizations of all *bona fide* immobile spots remained within that radius during their whole trajectories.

We then determined the fraction of trajectories that stay within a radius of 125 nm for δOR, κOR and μOR and would therefore be classified "immobile" by the above criterion. Hereby, we limited the trajectory lengths to a certain number of frames, to see the impact of shorter trajectory lengths (Suppl. Fig. 7). We found that when the trajectories were short, the apparently immobile fraction was much higher than for long trajectories. This obviously must be the case, since with shorter time, the area covered by the diffusing receptors will be smaller.

However, the longer the trajectories, the lower was the chance that a mobile receptor remained within the radius of 125 nm. The limiting value of the apparently immobile fraction for long trajectories is close to the actual fraction of immobile receptors.

In the single-molecule experiment without PhotoGate, we found that the fraction of immobile ORs after 15 frames (450 ms) was in the range of 10–20% (δOR: 18.9%; κOR: 11.9%; μOR: 13.4%). In the PhotoGate experiment, the fractions were around 5% (δOR: 4.5%; κOR: 4.4%; μOR: 5.5%). Accordingly, we can consider the fraction of mobile receptors to be only of minor influence for the results of our study.

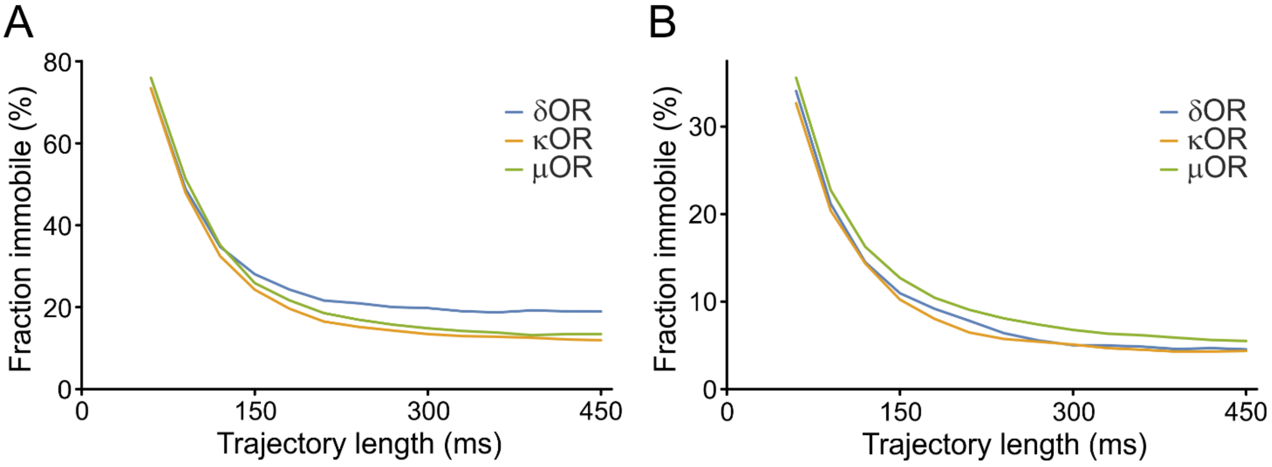


**Supplementary Figure 7: Fraction of immobile spots.** When limiting the trajectory length to a certain time, the apparent fraction of immobile spots is higher. In contrast, for long trajectories, it approaches the real value for the experiment **(A)** without PhotoGate and **(B)** with PhotoGate.

**Diffusion coefficient**

The diffusion coefficient of the molecules was determined by fitting a line to the function of mean squared displacement (MSD) over time. From a single movie, we selected all trajectories that contained 10 or more localizations. For these, we determined the MSD by calculating for each localization the displacement from the first point. A linear fit to this function (excluding the first point) yielded the 2D diffusion coefficient $d$ and localization uncertainty $\sigma$ from the equation

$$MSD(t) = 4Dt + 4\sigma^2$$

An example for a single low-expressing cell is shown in Suppl. Fig. 8. The average diffusion coefficient from all OR experiments was 0.065 ± 0.021 $\mu m^2/s$ (s.d.) (δOR: 0.061 ± 0.016 $\mu m^2/s$; κOR: 0.070 ± 0.023 $\mu m^2/s$; μOR: 0.064 ± 0.022 $\mu m^2/s$), and the average localization uncertainty was 35 ± 19 nm (s.d.). When excluding immobile spots, i.e. spots that remained within a circle of diameter 125 nm for their whole trajectory, the diffusion coefficient was 0.072 ± 0.021 $\mu m^2/s$ (s.d.) (δOR: 0.071 ± 0.018 $\mu m^2/s$; κOR: 0.076 ± 0.023 $\mu m^2/s$; μOR: 0.070 ± 0.023 $\mu m^2/s$), and the average localization uncertainty was 37 ± 20 nm (s.d.).

For high-expressing cells after using the PhotoGate, the average diffusion coefficient from all OR experiments was 0.086 ± 0.025 $\mu m^2/s$ (s.d.) (δOR: 0.087 ± 0.025 $\mu m^2/s$; κOR: 0.086 ± 0.022 $\mu m^2/s$; μOR: 0.084 ± 0.028 $\mu m^2/s$), and the average localization uncertainty was 41 ± 16 nm (s.d.). After excluding immobile spots, the diffusion coefficient was 0.090 ± 0.025 $\mu m^2/s$ (s.d.) (δOR: 0.092 ± 0.026 $\mu m^2/s$; κOR: 0.089 ± 0.021 $\mu m^2/s$; μOR: 0.089 ± 0.028 $\mu m^2/s$), and the average localization uncertainty was 41 ± 16 nm (s.d.)

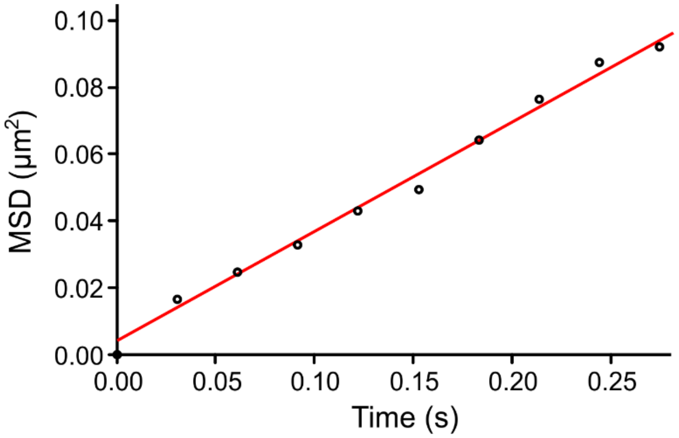


**Supplementary Figure 8: Mean squared displacement MSD from a κOR experiment.** 184 trajectories with a length of at least 10 localizations were evaluated. The diffusion coefficient was 0.082 $\mu m^2/s$ and the localization uncertainty 32 nm.

#### Supplementary Note 3

##### Determination of threshold distance for green and red to be considered co-localized

For the determination of green/red co-localization, a threshold distance is required. We therefore selected 20 *bona fide* yellow spots from the dually labeled SNAP549-PDGFRTM-GFP (spots where we determined visually by their co-movement that they were carrying both labels), which gave us a total of 808 localizations for each color. As an arbitrary criterion, we postulated that 95% of the localizations should be considered co-localized when applying the respective threshold. For a threshold distance of 150 nm, 76.4% of localizations were co-localized, for 200 nm, the fraction was 91.3%, and for 250nm, it was 97.4% (Suppl. Fig. 9). The value of 95% was reached with a threshold distance of 213 nm.

The relatively large threshold value can be explained by mainly two factors: (i) the lack of correction for chromatic aberrations of the Olympus 100x/NA1.70 objective. Therefore, different parts of the image display a different offset between green and red image. Since the chromatic aberration also depends on the coverslip thickness (which varies slightly due to production and repetitive cleaning in KOH), it is difficult to account for the shift using a stretching operation; (ii) when using alternating excitation, there is some diffusion of the molecules between the acquisition of the green and the red frame. With a diffusion coefficient of 0.08 $\mu m^2/s$ and a frame time of 30 ms, the displacement during one frame is about 100 nm.

Also with a smaller threshold distance of 200 nm or a larger threshold of 250 nm, all results obtained in our study were similar, and the conclusions were the same.

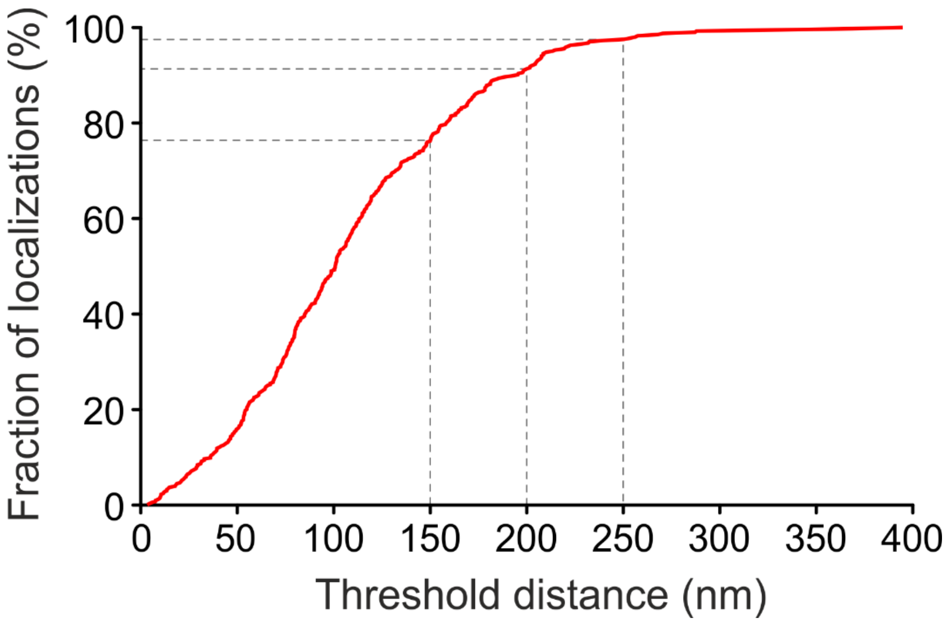


**Supplementary Figure 9: Determination of R/G threshold for co-localization.** From 808 localizations from 20 *bona fide* yellow spots, the fraction considered co-localized (red line) was determined in dependence of the threshold distance. Dashed gray lines indicate the co-localized fractions obtained with threshold values of 150, 200, and 250 nm.

#### Supplementary Note 4

##### Model to calculate dimer fraction from yellow fraction

The model takes into account the coincidental overlap of green and red florescent receptors, the formation of dimers from two green- or two red-labeled subunits, and the fractions of unlabeled SNAP-tag and non-fluorescent GFP. By assuming values for the total number of receptor subunits $N_0$, the fraction of total receptor subunits in monomers $f_m$, the fraction of GFP-labeled receptor subunits $f_G$, together with the known fractions of fluorescent GFP $p_g$ and labeled SNAP-tags $p_r$, and the ratio of spot area $A_s$ (spot radius $r_{spot}$) to area $A_0$ that all spots cover (i.e. the evaluated area of the membrane), we established a formula to estimate the values $N_{g-spot}$, $N_{r-spot}$, and $N_{y-spot}$ for the observed numbers of green, red and yellow spots.

The numbers of GFP- and SNAP-labeled monomers $N_{mG}$ and $N_{mS}$, and dimers carrying two GFP, two SNAP, or one of each $N_{dGG}$, $N_{dSS}$, $N_{dGS}$ are:

$$N_{mG} = N_0 \cdot f_G \cdot f_m$$

$$N_{mS} = N_0 \cdot (1 - f_G) \cdot f_m$$

$$N_{dGG} = \tfrac{1}{2} \cdot N_0 \cdot {f_G}^2 \cdot (1 - f_m)$$

$$N_{dSS} = \tfrac{1}{2} \cdot N_0 \cdot (1 - f_G)^2 \cdot (1 - f_m)$$

$$N_{dGS} = \tfrac{1}{2} \cdot N_0 \cdot 2 \cdot f_G \cdot (1 - f_G) \cdot (1 - f_m)$$

Due to non-fluorescent GFP and unlabeled SNAP-tags, some of these can appear dark, and dimers with GFP and SNAP can appear as yellow, green, red, or dark:

$$N_{mG\to g} = N_{mG} \cdot p_g$$

$$N_{mG\to d} = N_{mG} \cdot \left(1 - p_g\right)$$

$$N_{mS\to r} = N_{mS} \cdot p_r$$

$$N_{mS\to d} = N_{mS} \cdot (1 - p_r)$$

$$N_{dGG\to g} = N_{dGG} \cdot \left({p_g}^2 + 2 \cdot p_g \cdot \left(1 - p_g\right)\right)$$

$$N_{dGG\to d} = N_{dGG} \cdot \left(1 - p_g\right)^2$$

$$N_{dSS\to r} = N_{dSS} \cdot \left({p_r}^2 + 2 \cdot p_r \cdot (1 - p_r)\right)$$

$$N_{dSS\to d} = N_{dSS} \cdot (1 - p_r)^2$$

$$N_{dGS\to y} = N_{dGS} \cdot p_g \cdot p_r$$

$$N_{dGS\to g} = N_{dGS} \cdot p_g \cdot (1 - p_r)$$

$$N_{dGS\to r} = N_{dGS} \cdot \left(1 - p_g\right) \cdot p_r$$

$$N_{dGS\to d} = N_{dGS} \cdot \left(1 - p_g\right) \cdot (1 - p_r)$$

The numbers of yellow, green, red and dark appearing receptors are the sums:

$$N_y = N_{dGS\to y}$$

$$N_g = N_{mG\to g} + N_{dGG\to g} + N_{dGS\to g}$$

$N_r = N_{mS\to r} + N_{dSS\to r} + N_{dGS\to r}$

$N_d = N_{mG\to d} + N_{mS\to d} + N_{dGG\to d} + N_{dSS\to d} + N_{dGS\to d}$

Finally, we make an approximation to correct for the random co-localization due to close vicinity of red and green receptors. Then their emission overlaps spatially, resulting in appearance of yellow spots. A green receptor will appear as yellow spot when it happens to lie within the 213 nm radius of a red receptor. Therefore, the number of green and red spots is smaller than the number of receptors labeled in these colors, and the number of yellow spots is bigger. The chance for a single green receptor to not lie on a particular red receptor is $1 - A_s/A_0$, where $A_s = \pi \cdot r_{spot}{}^2$ is the area of the spot and $A_0$ is the total area the counted spots cover. The chance for a green receptor to lie on any of the red receptors is then:

$p_{rand} = 1 - (1 - A_s/A_0)^{N_r}$

and the number of yellow spots appearing from green and red overlap is:

$N_{y-rand} = N_g \cdot p_{rand}$

And the final counts of green, red and yellow spots are:

$N_{g-spot} = N_g - N_{y-rand}$

$N_{r-spot} = N_r - N_{y-rand}$

$N_{y-spot} = N_y + N_{y-rand}$

Estimates for the total number of receptor subunits $N_0$, the fraction of total receptor subunits in monomers $f_m$, and the fraction of GFP-labeled receptor subunits $f_G$ can be obtained by using them as fit parameters in a least squares fit of $N_{g-spot}$, $N_{r-spot}$, and $N_{y-spot}$ to the observed spot counts.

#### Supplementary Note 5

##### Model to calculate fluorescent GFP and labeled SNAP-tag fractions

The fractions of fluorescent GFP $p_g$ and labeled SNAP-tags $p_r$ can be obtained from the dually labeled SNAP549-PDGFRTM-GFP by counting the numbers $N_{g-spot}$, $N_{r-spot}$, and $N_{y-spot}$ of green, red and yellow labeled spots. The coincidental overlap of red and green receptors to form a yellow spot needs to be considered. The formulas from Suppl. Note 4 can be simplified because there are only receptors carrying both a GFP ad a SNAP-tag.

With the molecule number $N_0$, $p_g$ and, $p_r$ we directly get:

$$N_y = N_0 \cdot p_g \cdot p_r$$

$$N_g = N_0 \cdot p_g \cdot (1 - p_r)$$

$$N_r = N_0 \cdot \left(1 - p_g\right) \cdot p_r$$

The rest of the equations remains as in Suppl. Note 4:

$$p_{rand} = 1 - (1 - A_s/A_0)^{N_r}$$

$$N_{y-rand} = N_g \cdot p_{rand}$$

$$N_{g-spot} = N_g - N_{y-rand}$$

$$N_{r-spot} = N_r - N_{y-rand}$$

$$N_{y-spot} = N_y + N_{y-rand}$$

Estimates for the total number of receptor subunits $N_0$, the fraction of fluorescent GFP $p_g$, and the fraction of labeled SNAP-tags $p_r$ can be obtained by using them as fit parameters in a least squares fit of $N_{g-spot}$, $N_{r-spot}$, and $N_{y-spot}$ to the observed spot counts.

### Supplementary Note 6

##### Fraction of receptor subunits in dimers in dependence on total subunit density

The fraction $f_d$ of receptor subunits in dimers is

$$f_d = \frac{2[D]}{2[D]+[M]},$$

where $[M]$ and $[D]$ are the concentrations of monomers and dimers. The dissociation constant $k_d$ is given by

$$k_d = \frac{[M]^2}{[D]}$$

and the receptor subunit density $[R]$ is

$$[R] = 2[D] + [M].$$

Combining the three equations to eliminate $[M]$ and $[D]$ yields the quadratic equation

$$f_d{}^2 + (-2 - k_d/2[R])\, f_d + 1 = 0,$$

which has only one solution satisfying $0 \leq f_d \leq 1$, which is

$$f_d = 1 + \left(k_d - \sqrt{k_d(k_d + 8[R])}\right)/4[R].$$

#### Supplementary Note 7

##### Fitting of dissociation curve to κOR yellow spot fraction

Experiments for the κOR yielded green, red and yellow spot numbers $N_{g-spot}$, $N_{r-spot}$, and $N_{y-spot}$ in the (low-density) imaging area at pre-bleaching different receptor subunit densities $[R]$. It is important to note that this pre-bleaching subunit density $[R]$ and the density of (visible) receptor subunits in the imaging area $N_0/A_0$ are not the same; one is the density of the receptors before photobleaching, which determines the fraction of dimers and lies in the range of 10–150 / µm$^2$; the other is the density after photobleaching and partial re-population and is in the range of 0.2–2 / µm$^2$.

The green, red and yellow spot numbers $N_{g-spot}$, $N_{r-spot}$, and $N_{y-spot}$ depend on the receptor subunit number $N_0$, the fraction of total receptor subunits in monomers $f_m$, the fraction of GFP-labeled receptor subunits $f_G$, the fraction of fluorescent GFP $p_g$ and labeled SNAP-tags $p_r$, and the imaged/evaluated area $A_s$. However, the latter three of them are known for each experiment and therefore are not variables in the fitting procedure. The dependence of $N_{g-spot}$, $N_{r-spot}$, and $N_{y-spot}$ on the other parameters, as derived in Suppl. Note 4, can be described by a function

$$\{N_{g-spot}, N_{r-spot}, N_{y-spot}\} = g_{A_s}(N_0, f_m, f_g)$$

Since we make the assumption that the dimers do not dissociate during the 20 s between photobleaching and imaging, the fraction of subunits in monomers $f_m$ during imaging and the fraction of subunits in dimers $f_d$ before photobleaching (which in turn depends on the density $[R]$ and $k_d$) are related by

$$f_m = 1 - f_d([R], k_d)$$

$N_0$, $f_m$, $f_G$, $A_s$, and $[R]$ vary for each experiment $i = 1 \dots n$ and are therefore termed $N_0^i$, $f_m^i$, $f_g^i$, $A_s^i$, and $[R]^i$ in the following (not to be confused with an exponent). The same applies for $N_{g-spot}$, $N_{r-spot}$, and $N_{y-spot}$.

For finding the parameter $k_d$ that yields the best fit of the prediction to the observed values, we minimized the sum

$$\sum_{i=1\dots n} \left(\{N_{g-spot}^{i,pred}, N_{r-spot}^{i,pred}, N_{y-spot}^{i,pred}\} - \{N_{g-spot}^{i,obs}, N_{r-spot}^{i,obs}, N_{y-spot}^{i,obs}\}\right)^2 =$$

$$\sum_{i=1\dots n} \left(g_{A_s^i}\left(N_0^i, 1 - f_d\left([R]^i, k_d\right), f_g^i\right) - \{N_{g-spot}^{i,obs}, N_{r-spot}^{i,obs}, N_{y-spot}^{i,obs}\}\right)^2$$

by variation of all the $N_0^i$ and $f_g^i$ and the common $k_d$.

The 68% confidence interval was determined by the approach outlined in: Kemmer G. and Keller S. Nonlinear least-squares data fitting in Excel spreadsheets. *Nat. Protoc.* **5**: 267–281 (2010). Basically, $k_d$ was fixed to a value different than the one from the best fit, and fitting was repeated with the remaining free parameters. The confidence interval started and ended at the two values of $k_d$ where the sum of the squared residuals (SSR) reached 116.4% of the SSR from the best fit. The value 116.4% was determined from the Fisher distribution as exemplified in the reference cited above, with the number of adjustable parameters *M* = 3 and the number of data points *N* = 54 (3 numbers for each of the 18 experiments).
